# Interactions between inbreeding, fitness and the bacterial microbiome in *Aedes aegypti* mosquitoes

**DOI:** 10.64898/2026.03.14.711767

**Authors:** Perran A. Ross, Nadieh de Jonge, Qiong Yang, Véronique Paris, Torsten N. Kristensen, Ary A. Hoffmann

## Abstract

Laboratory and field populations of insects can experience a decline in fitness and loss of genetic diversity due to inbreeding depression and genetic drift, respectively. Matings among related individuals and small population size may also influence insect host microbiomes with consequences for fitness. In the dengue vector mosquito, *Aedes aegypti*, the bacterial microbiome is largely environmentally determined but recent studies have also revealed host genetic components. We generated a panel of 55 inbred lines from either of two founding outbred populations of *Ae. aegypti* to test for associations between life history traits, inbreeding, allelic diversity and microbiome composition using ddRADseq and bacterial 16S rRNA gene sequencing on pools of mosquitoes. Effects of inbreeding were diverse with severe composite fitness costs in many lines but minimal costs in others despite similar low levels of genetic diversity. We found no strong relationship between major life history traits across inbred lines, suggesting that any costs due to inbreeding were trait specific. Bacterial microbiome analysis of pooled samples from a subset of populations revealed common microbes across populations, with *Elizabethkingia*, *Aeromonas* and *Ralstonia* being the most abundant. Despite bacterial composition varying widely, there was no clear relationship between microbiome composition and fitness or population origin. However, there were several significant positive correlations between the relative abundance of different microbial taxa across lines. Our results demonstrate diverse impacts of inbreeding on fitness of mosquito populations but with limited impacts on the microbiome.

## Introduction

The *Aedes aegypti* (Diptera: Culicidae) mosquito is an invasive species that has spread rapidly throughout tropical, subtropical and even temperate regions within the last century and is responsible for hundreds of millions of arbovirus transmission events and billions of dollars in economic costs annually (Roiz et al., 2024). *Aedes aegypti* populations can be readily maintained in captivity and are widely used in research as a model system (Matthews and Vosshall, 2020). As one tool for mitigating the spread of pathogen transmission, *Aedes* mosquitoes are mass-reared in the laboratory and factory for release in biological control programs to suppress resident mosquito populations (Benedict, 2021, Dobson, 2021, Lim et al., 2024) or transform them to be refractory to pathogen transmission (Wang et al., 2024, Hoffmann et al., 2024, Gesto et al., 2021). These strategies are facilitated when mosquitoes are raised in artificial conditions to be fit and competitive under natural conditions (Nguyen et al., 2015, Garcia et al., 2020), thus understanding factors that influence mosquito fitness is important from an applied perspective.

Insect populations reared in artificial conditions can experience a decline in their future fitness under natural conditions for several reasons (Hoffmann and Ross, 2018, Lainhart et al., 2015, Azrag et al., 2016). When small populations are maintained in research settings, there may be a loss of fitness due to inbreeding depression, which is expected to lead to a general decline in overall fitness. In addition, in small populations, genetic drift can lead to a loss of alleles that might be favored under natural conditions. Even if laboratory populations are derived from many founders and kept at high population sizes, inbreeding and loss of genetic variation is often an unintended consequence of laboratory rearing (Briscoe et al., 1992, Hansen et al., 2025). In *Ae. aegypti*, inbreeding caused by low effective population size (Ne) can lead to severe fitness costs across a wide range of traits (Ross et al., 2019a). These issues are not expected to influence the fitness of factory stocks typically reared at a population size of many thousands to meet release targets (Zhang et al., 2018, Anders et al., 2025); however in establishing new factory lines, the stocks can be exposed to serial drift and bottleneck events as well as selection. Where experiments are being undertaken on long established lines, such as often used in pesticide testing (Hougard et al., 2003, Mosqueira et al., 2010), the situation is further compounded.

There is an increasing appreciation that the fitness of insect populations can be partly driven by the microbiome (Zheng et al., 2023). Recent work in *Drosophila melanogaster* has found that populations that have experienced severe genetic bottlenecks have reduced microbiome diversity, both of which contribute to decreased fitness in lines with the lowest genome-wide genetic variation (Ørsted et al., 2022). In *Aedes* mosquitoes, the bacterial microbiome is important for normal development, with axenic mosquitoes lacking a microbiome failing to develop or experiencing significant fitness costs (Correa et al., 2018, Coon et al., 2016). However, fitness can be partially or completely restored through the addition of specific microbes (Coon et al., 2016, Kriefall et al., 2024, Roman et al., 2024, Díaz et al., 2025). Mosquito larval habitat assessments have also provided links between microbiome composition and the productivity of larval habitats (Zhao et al., 2025). While mosquitoes lack obligate endosymbionts, facultative endosymbionts including *Wolbachia* can occur in mosquitoes and also have substantial effects on fitness (Ross et al., 2019b). It has been suggested that some species of mosquitoes carry a “core” microbiome (David et al., 2016, Guegan et al., 2018), with common microbes present in mosquitoes across different environments, but there is likely functional redundancy among bacterial groups (Rojas-Guerrero et al., 2024). Microbes present within mosquitoes can also affect their capacity to transmit pathogens (Cansado-Utrilla et al., 2021), including both environmentally acquired microbes and heritable endosymbionts (Herren et al., 2020, Moreira et al., 2009, Zhang et al., 2024, Wu et al., 2019).

Mosquito gut microbes are largely environmentally acquired, with their composition influenced by larval environment (MacLeod et al., 2021, Dickson et al., 2017, Zouache et al., 2022, Kriefall et al., 2024, Schwing et al., 2024) and conditions during adulthood including mating and blood and sugar feeding (Muturi et al., 2019, Chen et al., 2020, Díaz et al., 2021, Salgado et al., 2024). For instance, mosquito microbiomes differ fundamentally between lab and field populations (Didion et al., 2021, Hegde et al., 2018, LaReau et al., 2023, David et al., 2016). However, there are ways in which gut bacteria could be passed across mosquito generations, such as transfer of microbes from females to the larval environment through oviposition (Mosquera et al., 2023). While the environmental contribution to the mosquito microbiome is well established, there is a relatively limited understanding of genetic factors. Some studies have found that populations of *Ae. aegypti* larvae from distinct geographic locations tend to have the same gut microbiota when reared in a common environment, suggesting that the microbiome is entirely environmentally driven (Dickson et al., 2018, Scolari et al., 2019). Others identify differences between mosquito populations that persist in common environments (Accoti et al., 2023, Brettell et al., 2025, Kozlova et al., 2021, Short et al., 2017). This suggests the presence of a genetic component and potentially some host control over microbiome composition.

In earlier work (Ross et al., 2019a), we compared the fitness of *Ae. aegypti* populations at different levels of inbreeding ranging from complete outbreeding to consecutive generations of sib mating. The negative effects of inbreeding reflected the expected increased homozygosity by descent when contrasted to laboratory adaptation in some larger populations. Here we extend this work to consider the microbiome component by generating a panel of 55 inbred *Ae. aegypti* lines derived from two outbred populations to build up multiple effects of expression of recessive deleterious alleles and random allele fixation which was used to investigate any potential interactions with the microbiome. Given the importance of microbiomes for mosquito fitness and evidence for a heritable component, we hypothesized that the very low effective population size with expectant inbreeding depression may have strong impacts on the microbiome, potentially reducing microbiome diversity as seen in e.g. *Drosophila melanogaster* (Ørsted et al. 2022) with further negative impacts on fitness beyond those of the genetic bottlenecks of the host.

## Materials and methods

### Founding populations

We used two founding *Ae. aegypti* populations to establish inbred lines. Both populations were collected from Cairns, Australia (GPS: −16.921, 145.753) in 2014 and had been maintained at a census size of around 500 individuals for over 60 generations in the laboratory. Almost all *Ae. aegypti* in Cairns carry *Wolbachia* strain *w*Mel, an endosymbiont that was deliberately released to control dengue transmission by local mosquitoes (Hoffmann et al., 2011). Population A was collected from locations where releases of the *w*Mel *Wolbachia* strain had not occurred (Hoffmann et al., 2011, Ryan et al., 2019). Population B was collected from *Wolbachia* release zones, then *Wolbachia* was removed within the first 5 generations of laboratory rearing by treating larvae with 50 µg/L tetracycline hydrochloride in the larval rearing water and adults with 2 mg/mL tetracycline hydrochloride in a 10% sucrose solution for 2 consecutive generations. The complete absence of *Wolbachia* from Populations A and B was confirmed routinely using qPCR assays targeting *Wolbachia*- and mosquito-specific markers (Lee et al., 2012). *Wolbachia*-free mosquitoes were used because *Wolbachia* does not naturally occur in *Ae. aegypti* (Ross and Hoffmann, 2024) and it dominates the adult bacterial microbiome when present (Audsley et al., 2017). Mosquito populations were reared under laboratory conditions at 26°C with a 12:12 light:dark cycle according to methods described previously (Ross et al., 2017). Blood from a single human volunteer was used to feed all mosquito populations since establishment in the lab and for the duration of the study. Blood feeding on human volunteers was approved by the University of Melbourne Human Ethics committee (Project ID 28583).

### Inbreeding procedure

We established 70 inbred lines (46 from population A and 24 from population B) following a similar procedure to Ross et al. (2019a) by isolating mated, blood fed females from both populations in 70 mL specimen cups with a sandpaper oviposition strip (Ross et al., 2017). All offspring from a given female were placed together in a 1 L cage and allowed to mate. Adults were blood fed and two females per line were isolated for oviposition. If neither female produced offspring, additional females were isolated. Eggs from the remaining females were also collected and stored to be hatched if the line became extinct. The offspring from a single female was used to establish the next generation and this process was repeated for a total of 7 generations of sib mating. Since multiple paternity can occur at low frequencies in *Ae. aegypti* (Helinski et al., 2012, Richardson et al., 2015) this design resulted in either half sib or full sib crosses each generation. Populations A and B were maintained concurrently at a census size of ca. 500 individuals.

At G8 after 7 generations of sib mating, populations were amplified by collecting and breeding all offspring without further inbreeding, and they were maintained at a maximum census size of ca. 200 individuals. All 55 (15 of the original 70 went extinct) inbred lines and the two outbred populations were measured for life history traits (see description below) and sampled for ddRADseq and 16S rRNA gene sequencing at G10 (see below).

### Life history traits

We performed experiments at G8 and G10 to measure several life history traits of outbred and inbred lines. Our initial experiment at G8 was a pilot involving eight inbred lines to establish whether they differed in life history traits after the inbreeding procedure was completed. At G10, the experiment was repeated with all 55 remaining inbred lines and the two outbred populations.

We used identical methods in each experiment, with the exception that wing lengths were only measured at G10. Eggs (< 1 week old, stored at high relative humidity) were hatched in reverse osmosis (RO) water with a few grains of yeast. Within 6 hr of hatching, larvae were counted into 750 mL plastic trays filled with 500 mL of reverse osmosis (RO) water. We established up to 5 replicate trays with 50 larvae per tray for each mosquito line. Due to poor performance of some inbred lines, 7 lines at G10 and 3 lines at G20 had fewer than 250 larvae hatching and therefore fewer than 5 replicates of 50 larvae were established for these lines. Larvae were provided with food (Hikari Tropical Sinking Wafers, Kyorin Food, Himeji, Japan) *ad libitum* until pupation. Larval development time was measured for each sex by counting and sexing pupae twice daily in each tray. Survival to pupation was estimated by dividing the number of pupae that emerged by the initial number of larvae in each tray. Sex ratio was estimated at the pupal stage by dividing the number of male pupae by the total number of pupae. Due to logistical constraints, pupae from all replicate trays were added to a single container for adult emergence, with pupal mortality recorded across all replicates of a line by counting pupae that died or failed to eclose. Adults were released into 3 L cages and provided with water and a 10% sucrose solution. Ten males and 10 females from each line were stored in absolute ethanol and measured for their wing length according to methods described previously (Ross et al., 2017). Approximately 6 days after emergence, sucrose was removed for 24 hr whereafter females from each line were blood fed on the forearm of a single human volunteer. Up to 20 engorged females per line were isolated in 70 mL specimen cups for oviposition, though the level of replication varied due to mortality or failure to feed. Eggs were collected daily for up to one week then hatched three days post-collection. Fecundity was measured by counting the total number of eggs and their hatch proportion was measured by dividing the number of eggs with a clearly detached cap by the total number of eggs. We also calculated the proportion of females that blood fed but did not lay eggs in each line. We computed composite fitness for each line by multiplying the mean number of larvae per female by the proportion of larvae surviving to the adult stage. This value was then multiplied by the proportion of females laying eggs.

### ddRADseq

We used pooled double-digest RADseq (ddRADseq) to determine the effective population size (*N*e) of the two outbred populations and 55 inbred lines at G10 relative to the two founding populations at G0. We prepared pooled libraries following methods described previously for individuals (Rašić et al., 2014, Schmidt et al., 2018) and modified for pooled mosquitoes (Ross et al., 2019a). DNA was extracted from one pool of 20 male mosquito heads from each line using a Roche DNA Isolation Kit for Cells and Tissues (Roche, Pleasanton, CA, USA). DNA from each pool was quantified using a Qubit dsDNA HS Assay Kit and a Qubit 2.0 Fluorometer (Thermo Fisher Scientific, Life Technologies Holdings Pte Ltd, Singapore) and normalized.

We started with an initial digestion of 10 ng of genomic DNA, ten units each of MluCI and NlaIII restriction enzymes, NEB CutSmart buffer (New England Biolabs, Beverly, MA, USA), and water. Digestions were run for 3 h at 37 °C with no heat kill step, and the products were cleaned with Ampure XP™ paramagnetic beads (Beckman Coulter, Brea, CA, USA). Modified Illumina P1 and P2 adapters were ligated onto cleaned digestions overnight at 16 °C with 1000 units of T4 ligase (New England Biolabs, Beverly, MA, USA), followed by a 10-min heat-deactivation step at 65 °C. We performed size selection using a Pippin-Prep 2% gel cassette (Sage Sciences, Beverly, MA) to retain DNA fragments of 450–650 bp. The size selected libraries were amplified by PCR, using 1 μL of size-selected DNA, 5 μL of Phusion High Fidelity 2× Master mix (New England Biolabs, Beverly MA, USA) and 2 μL of 10 μM standard Illumina P1 and P2 primers. These were run for 12 PCR cycles, then cleaned and concentrated using 0.8x paramagnetic beads. Each ddRAD library contained 24 mosquito pools and was sequenced on a single sequencing lane using 2 × 100 bp paired-end reads.

### Data processing and genetic diversity estimates

We used the *Process_radtags* program in *Stacks* v2.0 (Catchen et al., 2013) to demultiplex sequence reads. Low-quality reads were discarded using a 15 bp sliding window if the average Phred score dropped below 20. We used *Bowtie2* v2.3.4.3 (Langmead and Salzberg, 2012) to align reads to the *Ae. aegypti* reference genome AaegL5.0 (Matthews et al., 2018). Alignments were converted to SAM format and sorted with SAMtools v1.7 (Li et al., 2009). We then used SAMtools to convert sorted files to the mpileup format, with each file containing one of the lines at G10 and its founding population at G0. These files were converted to sync format using the mpileup2sync.jar tool from PoPoolation2 (Kofler et al., 2011). We then estimated effective population size (*N*e) using the Nest R package v1.1.9 with two different methods (Jorde and Ryman, 2007, Jonas et al., 2016). Some lines returned NA values, likely due to insufficient SNPs or low allele frequency variance caused by strong inbreeding resulting in unreliable Ne estimation, and these were excluded from the Ne analyses.

As an additional measure of genetic diversity, we quantified the loss of founder variation between G0 and G10. Variants were called independently for each pooled ddRADseq sample using FreeBayes v1.3.2 (Garrison and Marth, 2012) under the --pooled-continuous model with --report-monomorphic, --exclude-unobserved-genotypes, and --standard-filters enabled. A minimum coverage threshold of 40× was required during variant calling. VCF files were processed using BCFtools v1.9 (Danecek et al., 2021). Sites with missing genotypes or mean depth <50× or >300× were removed. Downstream analyses were restricted to high-confidence sites passing these depth and genotype filters. For each line, we calculated the proportion of callable founder-variable sites that were fixed (reference or alternate). Callable sites were those with sufficient coverage to assign genotype state. Lines with fewer than 10 callable sites were excluded from analysis.

### 16S rRNA metabarcoding

At G10, 37 inbred lines and both outbred populations were sampled for 16S rRNA gene sequencing with 1-2 pools of 5 female mosquitoes (< 24 hr old, not fed sugar) per population. High-quality DNA extractions were submitted to Novogene (Novogene Co. Ltd, Hong Kong) for library construction using the universal 341F and 806R primers targeting the hypervariable V3-V4 region of the bacterial 16S rRNA gene. The raw paired-end Illumina sequencing data was processed using the AmpProc pipeline version 5.1 (GitHub - eyashiro/AmpProc: Illumina sequenced amplicon reads processing workflowGitHub - eyashiro/AmpProc: Illumina sequenced amplicon reads processing workflow), with the following settings: minimum amplicon length 200 bp, removal of 20 bp from both sides of the amplicon to strip primers, and clustering to Amplicon Sequencing Variants (ASV) and taxonomic assignment using SILVA 138.1.

### Microbiome analysis

Microbiome analysis was performed in the statistical software R (version 4.4.2), wrapped by RStudio. Analysis was performed using base R packages as well as ggplot2 (Wickham, 2016), ampvis2 (Andersen et al., 2018) and corrplot (Wei et al., 2017b). Alpha diversity was investigated using the observed richness (unique ASVs) and Shannon’s evenness metrics. Beta diversity was visualized using Principal Component Analysis (PCA) based on Hellinger-transformed abundance counts, as well as Principal Coordinate Analysis (PCoA) based on Robust Aitchison distances as implemented in the R package vegan (Oksanen et al., 2007). This approach is robust to sparsity and compositionality in microbiome data and is recommended for proportional comparisons of biodiversity (Gloor et al., 2017). In contrast, Hellinger transformation followed by Bray-Curtis distance can lead different conclusions about levels of biodiversity across treatments especially when using compositional datasets so caution is required in their interpretation (Fuschi et al., 2025, Seaton et al., 2023).

### Statistical analysis

To compare differences between lines and population origins across the life history traits, we ran ANOVAs using the car package (Fox et al., 2012) in R for each trait with population origin and line (nested within population origin) as factors. The threshold for significance was adjusted based on the number of traits measured using the Bonferroni correction. To assess correlations between traits, we first calculated the mean trait values from all replicate individuals or trays for each population. For egg hatch proportions which had a relatively high level of replication and where we detected significant deviations from normality, we calculated medians. Correlations between fitness (composite fitness and individual life history traits), effective population size (Ne(P), proportion fixed sites) and the microbiome data (number of unique ASVs, Shannon, and relative abundance of ASVs) were explored with Spearman’s correlations, using the cor.mtest R function with a significance level of 0.05 and a Bonferroni correction applied to account for the large number of pairwise comparisons undertaken.

## Results

### Diverse costs of inbreeding to life history traits despite similar and low Ne

After establishing 70 inbred lines through sib mating for 7 consecutive generations, 55 inbred populations remained at G8. The other 15 became extinct as none of the isolated females or backup eggs produced viable offspring. An initial assessment of eight inbred lines at G8 confirmed substantial differences between lines for fecundity, egg hatch proportion and survival to the pupal stage (Figure S1), with a significant effect of line for each of these three traits according to a one-way ANOVA (Table S1). We did not find a significant line effect for sex ratio or male development time and only a marginally significant line effect for female development time (Table S1).

Assessment of all 55 inbred lines and the two outbred populations revealed diverse effects of inbreeding on life history traits (Figure 1). Analyses of inbred lines found significant line (nested within origin) effects for all life history traits (Table S2). The outbred populations had higher composite fitness than the inbred lines (t-test, P < 0.001), however some inbred lines had higher fitness for certain traits such as fecundity and development time (Figure 1). Fitness differed between the inbred lines derived from Populations A and B, with significant effects of population origin for 5/8 traits that had within-line replication (Table S2), with a general tendency for greater inbreeding depression in Population B. For instance, inbred lines from Population B had lower fecundity (Figure 1A) and survival to the pupal stage on average but developed faster (Figure 1E-F) than lines from Population A.

**Figure 1.**
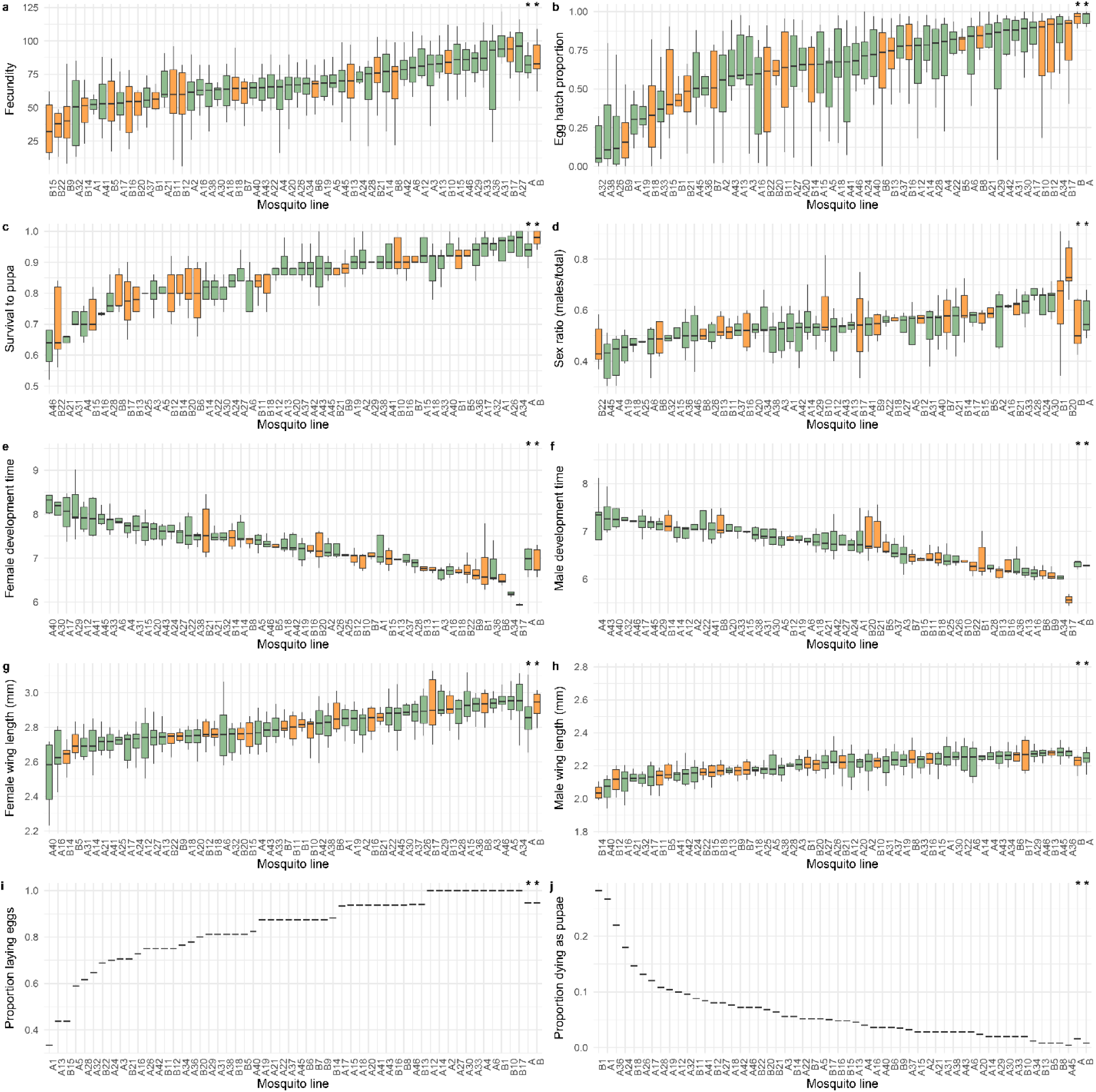
Life history traits of two outbred and 55 inbred lines of *Aedes aegypti* at G10. (A) Fecundity, (B) egg hatch, (C) survival to pupa, (D) sex ratio, (E) female development time, (F) male development time, (G) female wing length, (H) male wing length, (I) proportion of females laying eggs and (J) pupal mortality. The inbred lines are ranked by median trait value and are colored by the founding population (Population A in green and Population B in orange). Data for the two outbred populations are presented on the right-hand side and indicated by asterisks. Boxplots show medians and interquartile ranges. Panels (I) and (J) have a single value for each population as proportions were calculated from a pool of all replicates.

To confirm that inbred lines experienced significant bottlenecks, we performed ddRADseq on pooled samples from all populations at G10 and estimated genetic diversity relative to their founding population at G0. As expected, inbred lines had sharp reductions in Ne (range: 1.51-5.51 based on the Ne(P) method where there were fewer missing values, Table S3) compared to outbred populations A (Ne(P) = 77.99) and B (Ne(P) = 169.73). The inbred lines also had a relatively high proportion of variable sites in their founding population that became fixed (range: 0.867-1.000, Table S3) compared to outbred populations A (0.764) and B (0.789). Nevertheless, we did not find a significant positive correlation between genetic diversity and composite fitness among the inbred lines (Ne(JR): *rs* = −0.078, P = 0.603, Ne(P): *rs* = 0.224, P = 0.110, proportion fixed sites: *rs* = −0.168, P = 0.313, Figure S2). This could indicate that any differences in the loss of genetic variation due to sib mating is too similar across lines (or insufficient) to generate line specific effects on fitness.

To test whether line differences for fitness traits were associated, we performed pairwise correlations between all traits measured across the inbred lines (Figure 2a). We found strong correlations between male and female wing length (*rs* = 0.691, P < 0.001), and male and female development times (*rs* = 0.814, P < 0.001) across the lines, where lines with large females tended to have large males and lines where females were fast to develop tended to have males that were also fast to develop. These results point to a lack of sex-specific effect of line differences following inbreeding in these traits. There were also weak correlations between other related traits, including egg hatch proportion and the proportion of females laying eggs (*rs* = 0.337, P = 0.011), but these were not significant following Bonferroni correction (cf. Figure 2a with Figure S5a). There were no significant correlations between any other trait pairs across the lines (Figure 2a), suggesting that we failed to fix deleterious genes with effects across multiple traits in the inbred lines.

**Figure 2.**
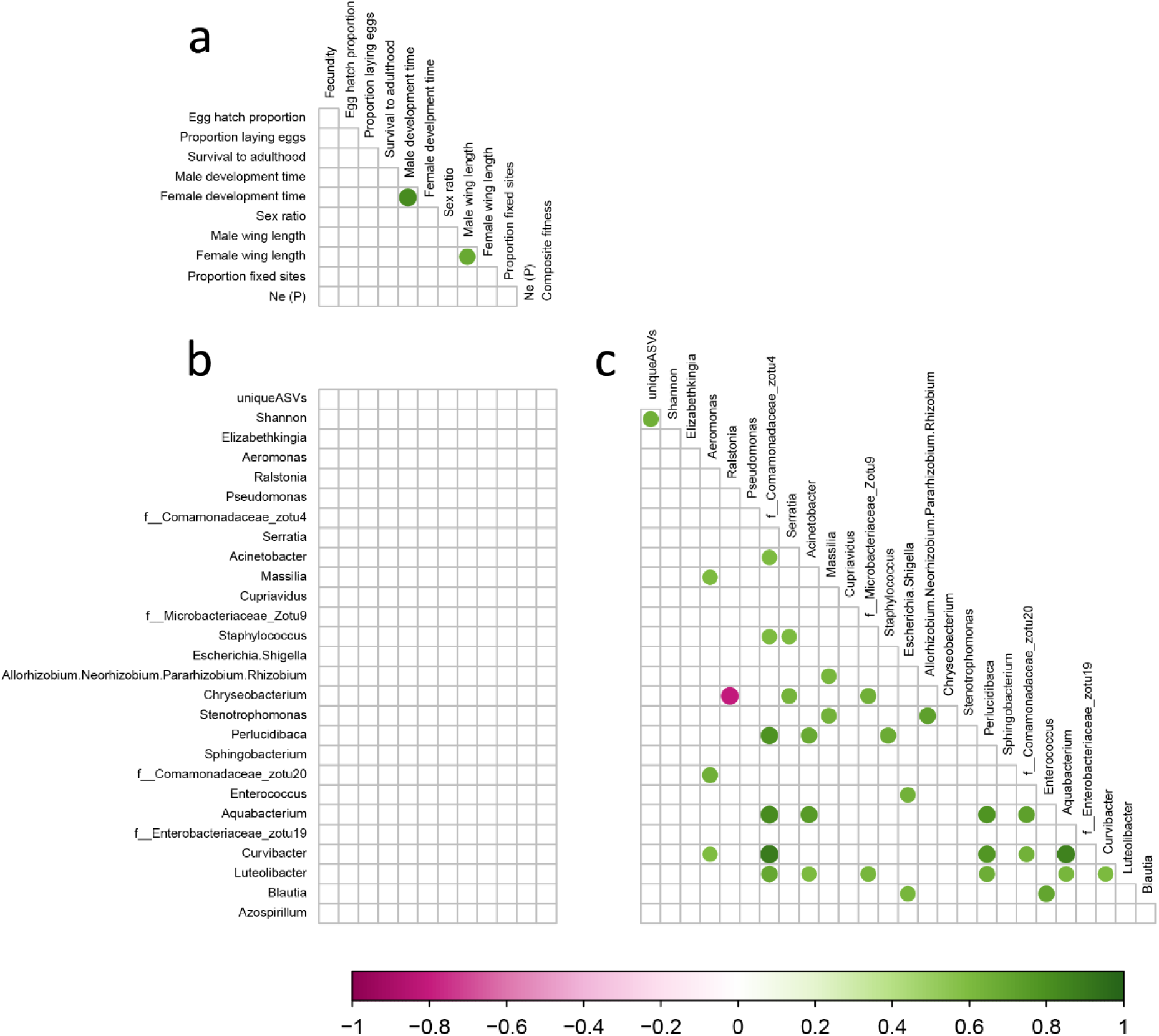
Spearman correlation matrix of life history traits, genetic diversity and microbiome composition across *Aedes aegypti* populations. Panel (A) shows correlations between the different life history traits as well as between measures of genetic diversity (Ne(P) and proportion fixed sites) and life history traits. Panel (B) shows correlations between life history traits (including composite fitness and genetic diversity) and microbiome traits (number of unique ASVs, Shannon index and the relative abundance of the 25 most common microbial taxa across all lines). Panel (C) shows correlations between the relative abundance of different microbes. Purple indicates a significant negative correlation while green indicates a positive correlation, and the circles are sized relative to the associated R^2^ value. Only significant correlations (P < 0.05 after Bonferroni correction) between tested variables are shown. See Figure S5 for an exploratory matrix where correlations have not been corrected for multiple comparisons.

### Diverse microbiome compositions of inbred lines

We collected samples of freshly emerged adults from populations at G10 and performed bacterial 16S rRNA gene sequencing on pools of mosquitoes. We obtained sequences from 37 inbred lines and both outbred populations. Bacterial microbiomes were dominated by a small number of genera with *Elizabethkingia*, *Aeromonas* and *Ralstonia* being the most abundant across lines (Figure S3). No endosymbionts (including *Wolbachia*) were detected in any sample. Out of 2354 ASVs across all lines, 59 had a relative abundance of > 0.01%, comprising 85% of all reads. In PCA plots of beta diversity (Figure 3) and a Robust Aitchison ordination (Figure S4), we found that outbred populations A and B clustered together, suggesting similarity in microbiome composition despite having different origins. The spatial distributions of the inbred lines derived from each outbred population also overlapped substantially, indicating a lack of population origin effect (Figure 3). There was no evidence for the inbred lines having reduced complexity or abundance in their microbiomes compared to the outbred populations as the two outbred lines had some of the lowest Shannon indices and lowest number of ASVs across all populations (Figure 4).

**Figure 3.**
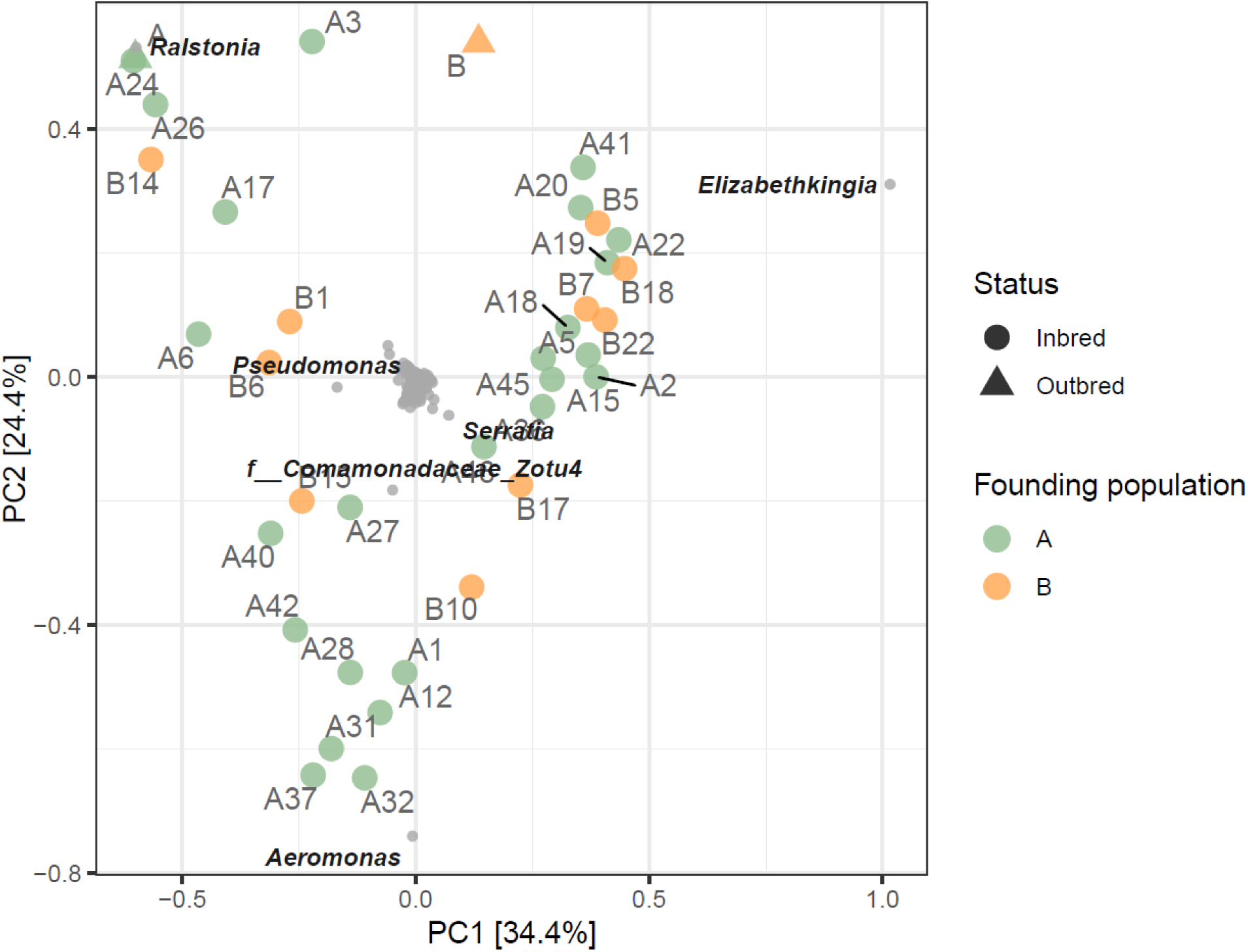
Principal component analysis of Hellinger-transformed bacterial relative abundance data. Samples are coloured by their founding population (A or B) and labelled by line number. The outbred lines are shown as triangles (population A is obscured in the top-left of the plot), while the inbred lines are shown as circles. Distances among samples reflect differences in overall microbiome composition. Bacterial species (ASVs) are displayed as species scores in the same ordination space, with their positions indicating the direction and strength of association with sample variation along the principal components. Species located closer to a group of samples are relatively more abundant in those lines. The six most influential species (based on contribution to the ordination axes) are labelled at the genus level or best available taxonomic resolution.

**Figure 4.**
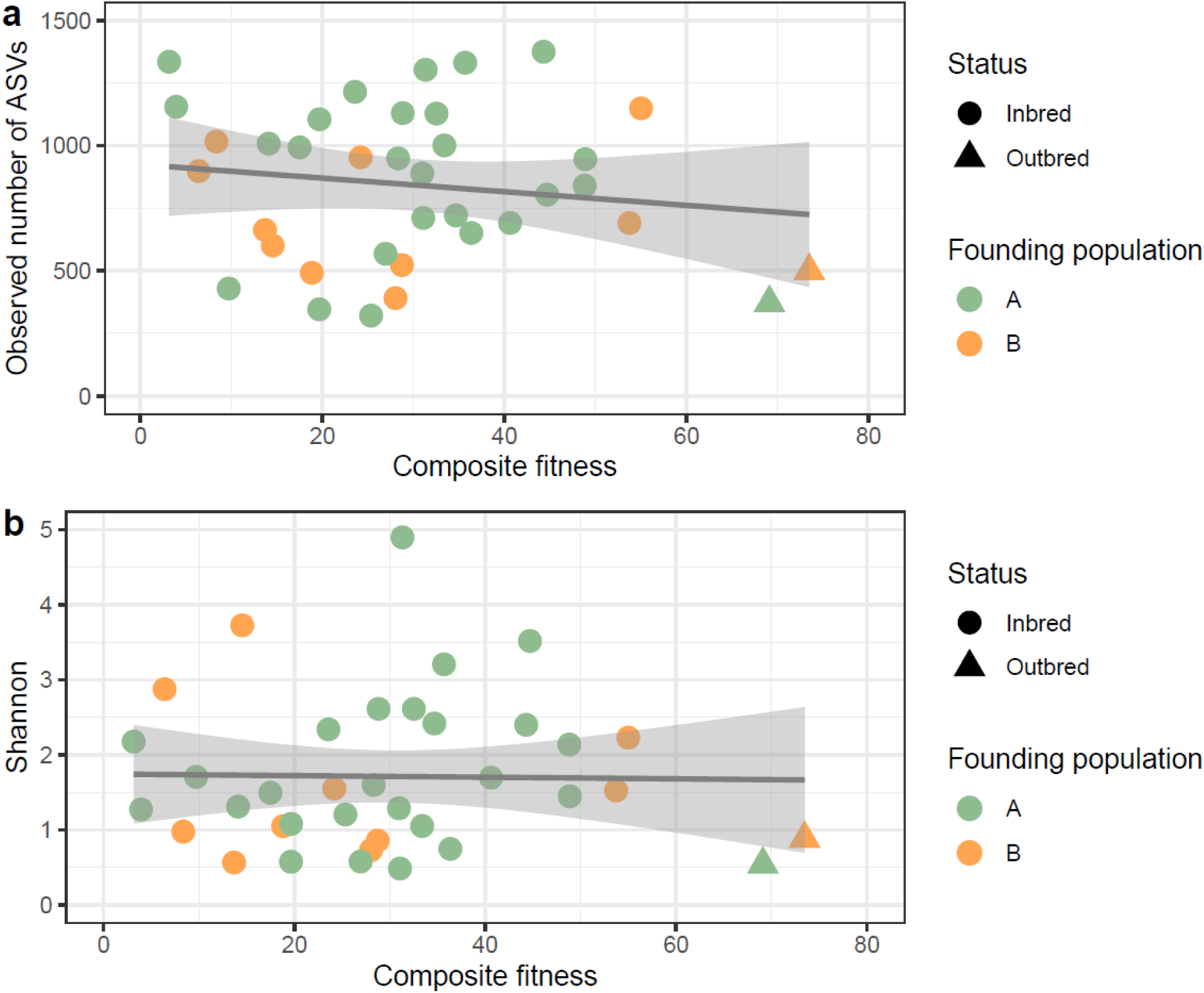
Association between bacterial 16S microbiome diversity and composite fitness. (A) Scatterplot of composite fitness and number of ASVs observed for each line. (B) Scatterplot of composite fitness and Shannon index values observed for each line. Points are colored by founding population (A or B) with triangle symbols indicating the two outbred lines. A linear regression line and 95% SE are drawn on each plot.

We then investigated correlations between the relative abundances of different microbial taxa across the lines (Figure 2c). We found several highly significant positive correlations between taxa (e.g. *Aeromonas* and *Massilia*) suggesting that these tend to co-occur with each other across the mosquito lines. In an exploratory analysis, we found that the relative abundance of *Ralstonia* was negatively correlated with the relative abundance of 10/24 of the most common microbial taxa with only one positive correlation (Figure S5). *Ralstonia* relative abundance also had a negative correlation with the number of unique ASVs (Figure S5). There may be antagonistic interactions between *Ralstonia* and the broader bacterial microbiome, where lines with a low abundance of *Ralstonia* tend to have more diverse microbiomes with a higher abundance of rarer taxa. However, these patterns should be interpreted with caution as correlations were not significant when the threshold was adjusted for the number of pairwise comparisons performed, except for a strong negative correlation between *Ralstonia* and *Chryseobacterium* relative abundance (*rs* = −0.803, P < 0.001, Figure 2c). In addition the 16S data we collected is proportional and caution is required when interpreting negative correlations (Gloor et al., 2017).

### Lack of association between line differences in fitness and the bacterial microbiome

The diversity of microbiome compositions across inbred lines allowed us to investigate potential interactions between life history traits and microbiome traits across lines. We found no significant correlation between composite fitness and the number of ASVs or Shannon indices (Figure 4). This was also the case when considering life history traits separately (Figure 2b), even when the threshold for significance was not adjusted (Figure S5b). We also did not find any strong associations between the relative abundance of specific microbes and any life history trait (Figure 2b), although there was a weak association between *Ralstonia* relative abundance and composite fitness (*rs* = 0.359, P = 0.026 – non-significant after Bonferroni correction) as well as several pairwise correlations between specific life history traits and microbial taxa (Figure S5).

## Discussion

Our experiments with a panel of inbred *Ae. aegypti* lines show that inbreeding has diverse effects on mosquito fitness despite substantial and relatively uniform decreases in genetic diversity. We also found large differences in bacterial microbiome composition across lines but no strong evidence for interactions between life history and microbiome characteristics. The results are broadly consistent with other data for *Ae. aegypti* demonstrating a stronger influence of environmental over genetic contributions to the microbiome (Accoti et al., 2023, Hyde et al., 2023), while contrasting with data from *Drosophila* showing that genetic bottlenecks substantially reduce microbiome diversity (Ørsted et al., 2022). As discussed below, this work has implications for fitness assays of mosquito populations performed under laboratory conditions and releases of mass-reared insects for pest and disease vector control.

The inbred populations of *Ae. aegypti* varied widely in their composite fitness despite consistently low Ne and high frequencies of fixed alleles. We previously reported a strong association between fitness and Ne in populations maintained at different census sizes (Ross et al., 2019a), but here we identify cases where the fitness of highly inbred lines are at similar levels to outbred populations. There were overall differences in fitness between inbred lines from the two origins which may reflect variation in the presence of deleterious recessive alleles in the two founding populations. We acknowledge that the costs of inbreeding identified in our study may be underestimated since we could only perform experiments with the lines that remained at G10. Fifteen of the 70 initial inbred lines became extinct and some of the remaining lines were derived from backups, so the remaining lines represent relatively fitter lineages. Consistent with previous studies (Ross et al., 2019a, DeRose and Roff, 1999, Bechsgaard et al., 2013), fertility-related traits were particularly susceptible to impacts of inbreeding depression, with egg hatch proportions in some inbred lines approaching zero. Although not formally tested, we observed that most inbred lines that became extinct did so through females either not laying eggs or through no eggs hatching, rather than as a consequence of mortality at the larval, pupal or adult stages.

With the large number of lines and traits measured for fitness, we could test for correlations between different traits across the lines. We expected to see associations between related traits such as body size and fecundity, for which there is a well-established relationship within populations of mosquitoes (Briegel, 1990) and in insects more generally (Honěk, 1993). Aside from strong relationships between male and female data for the same trait, no significant correlations between traits were observed across our panel of lines. This suggests that deleterious alleles affecting different traits are being fixed in the inbred lines.

Our assessment of mosquito microbiomes identified a core group of common microbes across the mosquito lines, with *Elizabethkingia*, *Aeromonas* or *Ralstonia* found at a high abundance in most lines. This aligns with other studies that identify a small group of common microbes across mosquito species despite their genetic divergence (Hyde et al., 2023). However, there was a diversity in the relative abundance of microbes across the lines despite being reared in a common environment and being derived from only two founding populations. The most abundant taxa have all been reported to colonize mosquitoes: *Elizabethkingia* (Chen et al., 2015), *Aeromonas* (Pidiyar et al., 2002), *Ralstonia* (Krajacich et al., 2018), *Pseudomonas* (Chavshin et al., 2015), *Serratia* (Mancini et al., 2018) and bacteria from the comamonadaceae family (Nilsson et al., 2018). We also found correlations between bacterial taxa across the lines, which suggests that some taxa covary across lines. Although most significant correlations were positive, we identified one strong and several weak negative correlations between *Ralstonia* and several other taxa which may reflect antagonistic interactions, where the presence of *Ralstonia* inhibits the colonization of other taxa (Hegde et al., 2018). Microbiome interactions in mosquitoes have been well-documented (Hegde et al., 2018), particularly for *Asaia* and *Wolbachia* which can mutually exclude the establishment and transmission of the other (Rossi et al., 2015, Hughes et al., 2014). However, since we sequenced pools of mosquitoes rather than individuals, we can only infer general patterns across lines rather than demonstrate interference between the two taxa within individual mosquitoes.

Few studies have explored links between inbreeding and microbiome diversity. One study in pigs identified decreased gut microbiome diversity in an inbred breed (Wei et al., 2020), while in aphids inbreeding can lead to a decline in the abundance of the obligate endosymbiont *Buchnera* (Matsuda et al., 2024). In contrast, one study of inbred and hybrid lines of maize showed that microbiomes were largely driven by non-genetic effects (Schultz et al., 2023). Based on experiments in *D. melanogaster* (Ørsted et al., 2022) and the fact that some common mosquito microbes can be transmitted across generations (Akhouayri et al., 2013, Mosquera et al., 2023), we hypothesized that genetic bottlenecks may constrain the mosquito microbiome. Despite our inbred lines having low and relatively uniform genetic diversity, there was a large spread of microbiome diversity across the lines and the outbred lines had some of the lowest microbiome diversity. Therefore there was no clear evidence that genetic bottlenecks led to microbiome bottlenecks in contrast to findings in *Drosophila*. Differences in microbiomes between lines are instead likely to reflect environmental contributions. Although all mosquito lines tested were reared in a common environment at the same time, we did not use any form of sterilization. The lines could therefore have been exposed to different microbes on the surface of egg papers, rearing containers or the insectary itself. Recent work shows that even subtle differences in their rearing conditions can have significant impacts on the mosquito microbiome (Brettell et al., 2025).

One of the main goals of our study was to test for a link between fitness and microbiome composition in our panel of inbred lines. Some of the common taxa identified in our populations have been already linked to fitness through experimental manipulation of the microbiome. For instance, some strains of *Aeromonas* (Wehbe et al., 2025) and *Serratia* (Wei et al., 2017a, Kozlova et al., 2021) have been described as pathogenic, while exposure to *Pseudomonas* spp. can also affect mosquito life history traits (Silva et al., 2024). We found limited evidence for associations between microbes and fitness, suggesting that any costs due to inbreeding likely reflect the increased expression of deleterious alleles, not due to an abundance of specific microbes. Given the diversity in microbiome composition between lines despite being reared in a common environment, it may prove challenging to identify associations with fitness under conventional rearing conditions. Gnotobiotic mosquitoes with a defined microbiome have recently been used to characterize the effects of single bacterial species on fitness (Díaz et al., 2025), and future work could extend this by generating panels of lines reared with diverse combinations of microbes or different larval diets to further establish links between fitness and the microbiome.

In summary, our study reinforces the need to consider the inbreeding status of mosquito lines used for research and release programs given the substantial fitness costs of inbreeding present in the inbred lines (Ross et al., 2019a). This is not so much an issue during rearing for mass release, but it becomes important when rearing lines are reinvigorated with new material from the field that aims to reduce the negative effects of laboratory adaptation. Our data do not point to specific components of the microbiome as further exacerbating the impacts of inbreeding in contrast to findings from some other species. However, this does not preclude the possibility of adding specific microbes during rearing to boost the fitness of mass-reared mosquitoes.

## Acknowledgements

We thank Meng-Jia Lau, Jessica Home, Xuefen Xu, Ashritha Dorai, Ella Yeatman, Mel Berran and Apeksha Warusawithana for assistance with line maintenance and technical assistance with the experiments. We thank Monica Stelmach and Nancy Endersby-Harshman for technical assistance with DNA extraction and sequencing. We also thank Joshua Thia, Thomas Schmidt and Moshe Jasper for bioinformatics assistance and advice.

## Data availability

Data from fitness experiments are available on Figshare at doi:10.26188/31403337. ddRADseq libraries and bacterial 16S rRNA gene sequencing data will be available upon acceptance of the manuscript.

## Supporting information

**Figure S1.**
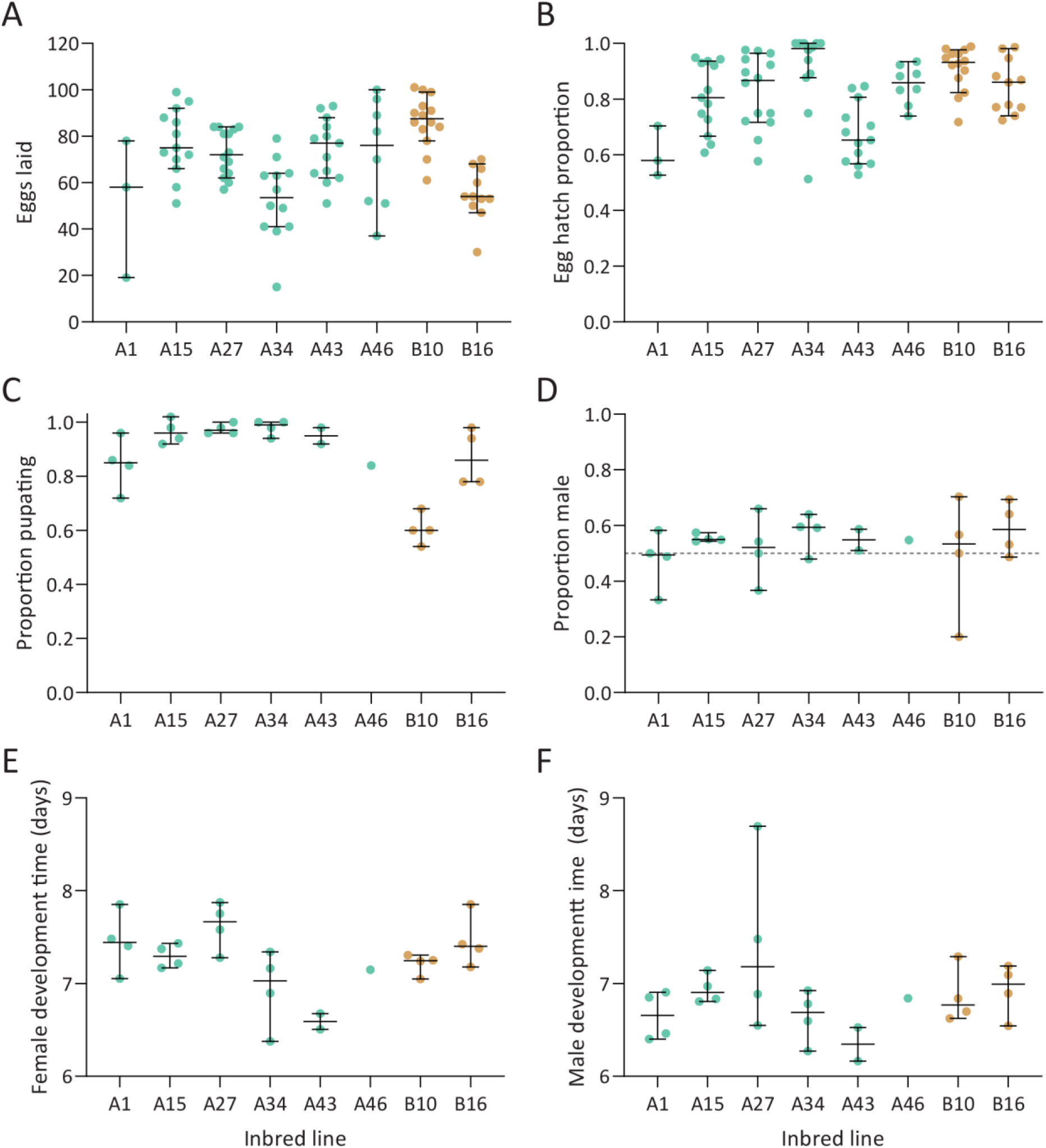
Life history traits of select inbred *Aedes aegypti* populations at G8. (A) Fecundity, (B) egg hatch, (C) survival to pupa, (D) sex ratio, (E) female development time and (F) male development time. Horizontal lines and error bars show medians and 95% confidence intervals with dots showing data from individual females (A-B) or replicate containers (C-F).

**Figure S2.**
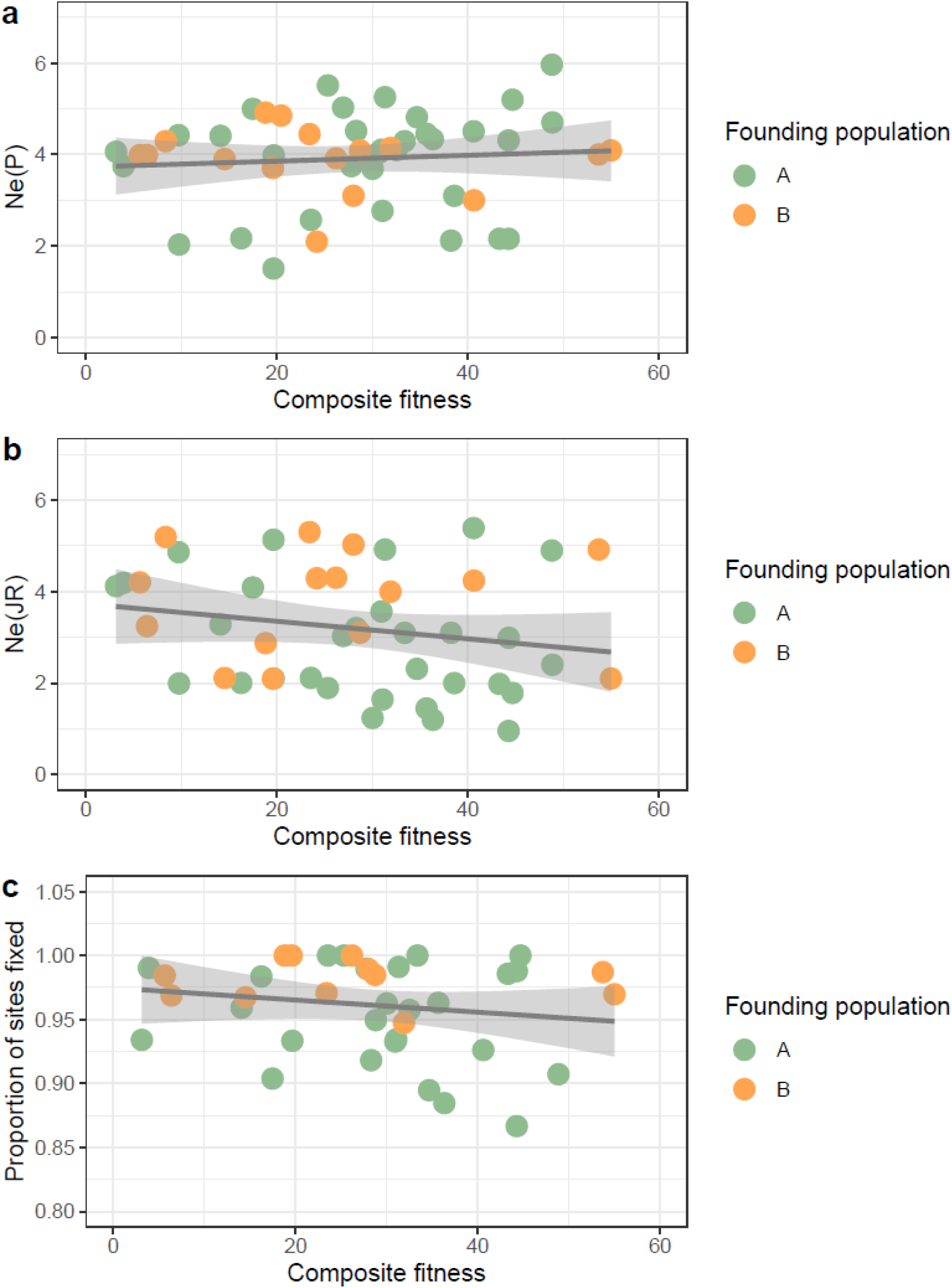
Regression between composite fitness and estimates of genetic diversity of inbred *Ae. aegypti* lines using the (a) Ne(P) and Ne(JR) methods for Ne and (c) the proportion of variable sites from the founding populations that became fixed. **Shaded areas represent 95% SE.**

**Figure S3.**
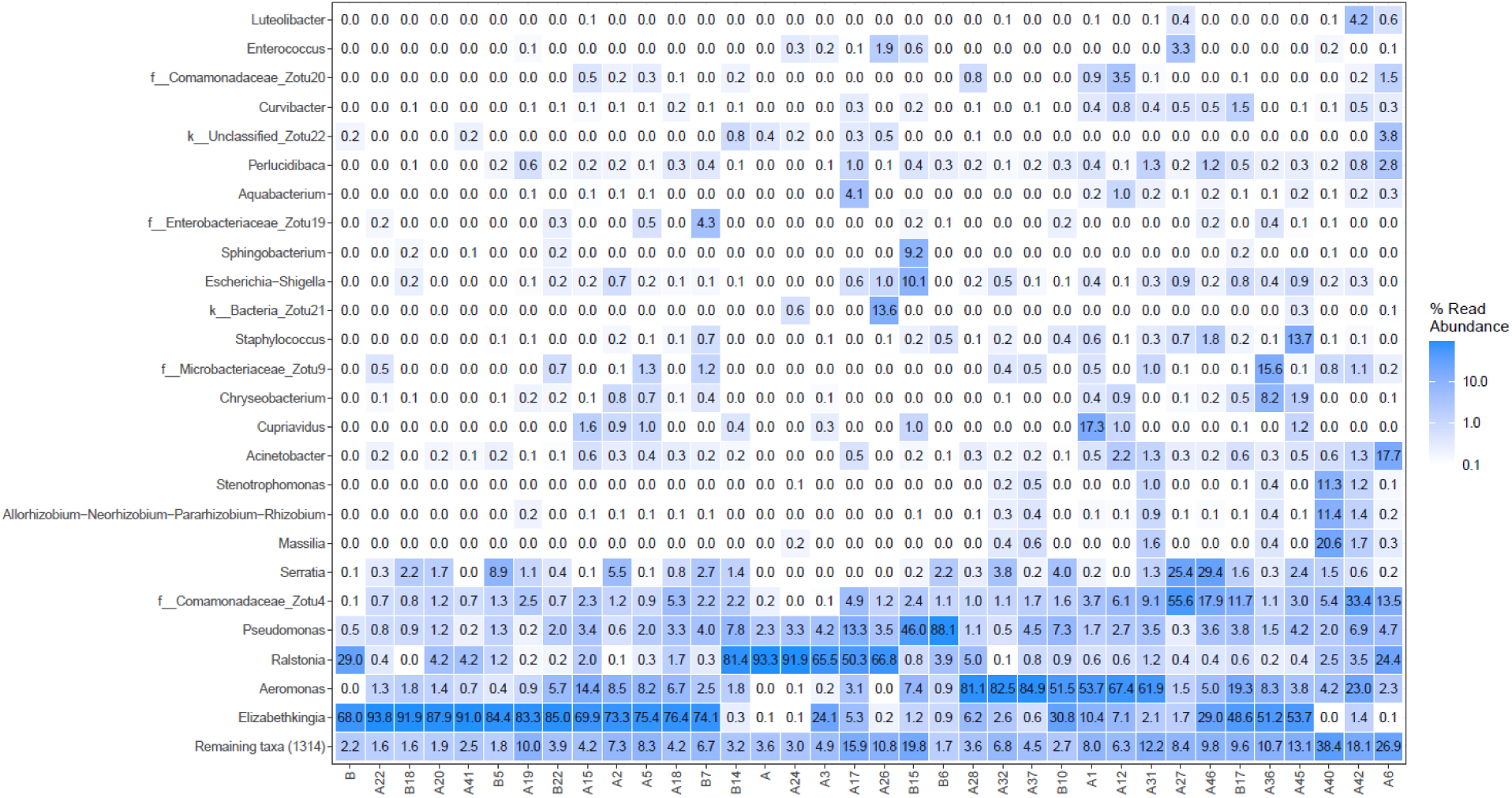
Relative read abundance of specific taxa for each *Ae. aegypti* line.

**Figure S4.**
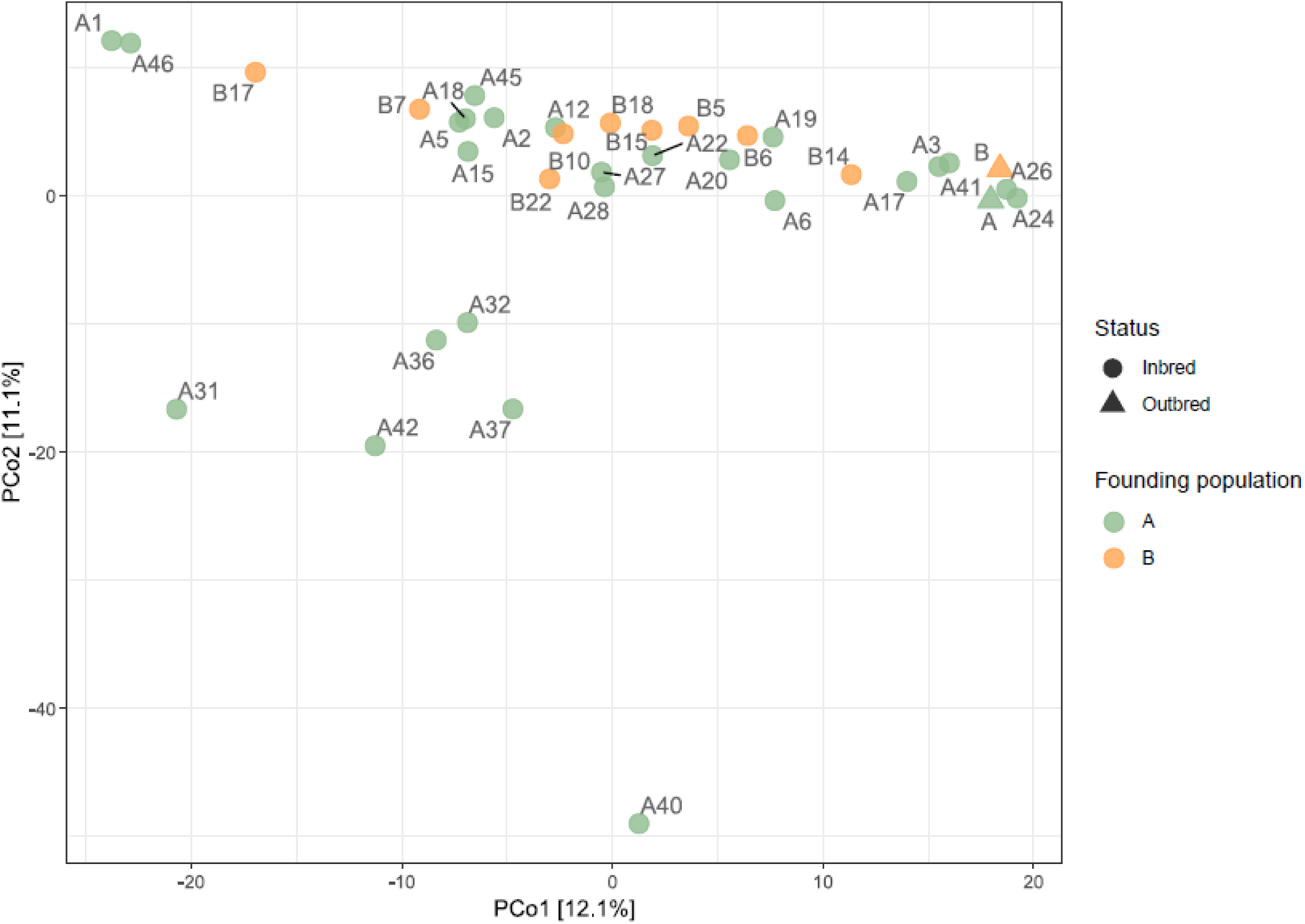
Robust Aitchison ordination of bacterial community composition across *Aedes aegypti* lines. Ordination was performed on robust Aitchison distances calculated based on ASV relative abundance data visualized by PCoA. Each point represents an individual line and is labelled by line ID.

**Figure S5.**
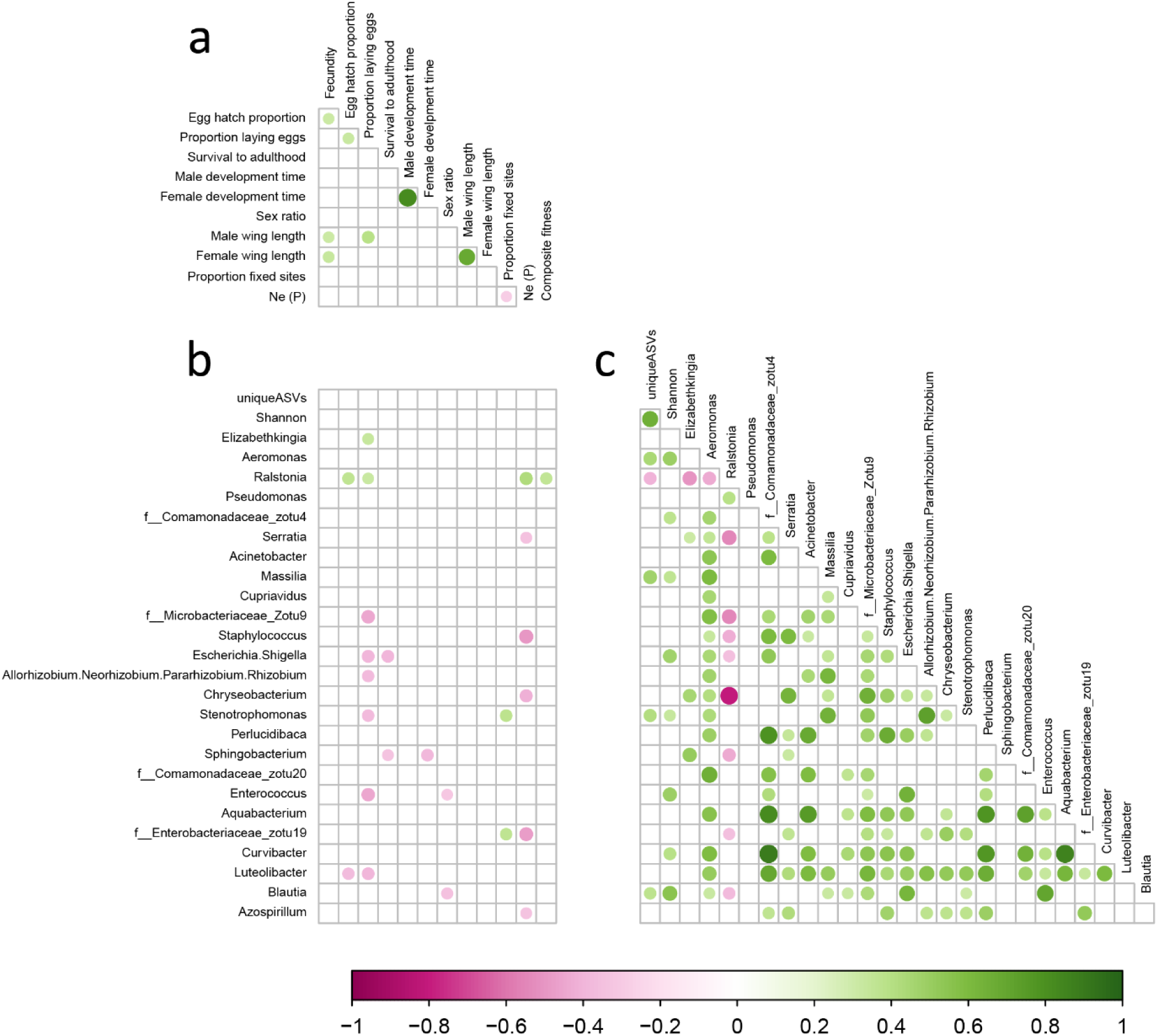
Spearman correlation matrix of life history traits, genetic diversity and microbiome composition across *Aedes aegypti* populations when not corrected for multiple comparisons. Panel (A) shows correlations between the different life history traits as well as between measures of genetic diversity (Ne(P) and proportion fixed sites) and life history traits. Panel (B) shows correlations between life history traits (including composite fitness and genetic diversity) and microbiome traits (number of unique ASVs, Shannon index and the relative abundance of the 25 most common microbial taxa). Panel (C) shows correlations between the relative abundance of different microbes. Purple indicates a significant negative correlation while green indicates a positive correlation, and the circles are sized relative to the associated R^2^ value. Only significant correlations (P < 0.05) are shown. These results are presented for exploratory purposes only and should be interpreted with caution given the large number of pairwise comparisons performed. Formal inference is based on the multiple-testing-corrected results shown in Figure 2.

**Table S1.** ANOVAs for life history traits of select inbred *Aedes aegypti* populations at G8.

| Source | Degrees of freedom | F | P value |
| --- | --- | --- | --- |
| **Fecundity** |  |  |  |
| Intercept | 1 | 1430.132 | < 0.001 |
| Inbred line | 7 | 7.739 | < 0.001 |
| Error | 80 |  |  |
| Total | 88 |  |  |
| **Egg hatch proportion** |  |  |  |
| Intercept | 1 | 3554.463 | < 0.001 |
| Inbred line | 7 | 7.433 | < 0.001 |
| Error | 80 |  |  |
| Total | 88 |  |  |
| **Survival to pupa** |  |  |  |
| Intercept | 1 | 3752.986 | < 0.001 |
| Inbred line | 7 | 13.801 | < 0.001 |
| Error | 19 |  |  |
| Total | 27 |  |  |
| **Sex ratio** |  |  |  |
| Intercept | 1 | 456.380 | < 0.001 |
| Inbred line | 7 | 0.452 | 0.857 |
| Error | 19 |  |  |
| Total | 27 |  |  |
| **Female development time** |  |  |  |
| Intercept | 1 | 15083.815 | < 0.001 |
| Inbred line | 7 | 4.159 | 0.006 |
| Error | 19 |  |  |
| Total | 27 |  |  |
| **Male development time** |  |  |  |
| Intercept | 1 | 5026.571 | < 0.001 |
| Inbred line | 7 | 1.517 | 0.221 |
| Error | 19 |  |  |
| Total | 27 |  |  |

**Table S2.** ANOVAs for life history traits of inbred *Ae. aegypti* populations at G10.

| Source | Sum of squares | Degrees of freedom | F | P value |
| --- | --- | --- | --- | --- |
| **Fecundity** |  |  |  |  |
| Origin | 14403.15 | 1 | 42.1958 | < 0.001 |
| Line (origin) | 103337.96 | 52 | 5.8220 | < 0.001 |
| Error | 214020.51 | 627 |  |  |
| **Egg hatch proportion** |  |  |  |  |
| Origin | 0.0108 | 1 | 0.1826 | 0.6693 |
| Line (origin) | 20.8108 | 52 | 6.7677 | < 0.001 |
| Error | 37.0776 | 627 |  |  |
| **Survival to pupa** |  |  |  |  |
| Origin | 0.0358 | 1 | 7.0281 | 0.0086 |
| Line (origin) | 1.5624 | 53 | 5.7873 | < 0.001 |
| Error | 1.0544 | 207 |  |  |
| **Sex ratio** |  |  |  |  |
| Origin | 0.0768 | 1 | 10.9829 | 0.0011 |
| Line (origin) | 0.9487 | 53 | 2.5596 | < 0.001 |
| Error | 1.4476 | 207 |  |  |
| **Female development time** |  |  |  |  |
| Origin | 13.5811 | 1 | 104.8124 | < 0.001 |
| Line (origin) | 46.1733 | 53 | 6.7234 | < 0.001 |
| Error | 26.8222 | 207 |  |  |
| **Male development time** |  |  |  |  |
| Origin | 7.0728 | 1 | 90.7918 | < 0.001 |
| Line (origin) | 30.2672 | 53 | 7.3308 | < 0.001 |
| Error | 16.1256 | 207 |  |  |
| **Female wing length** |  |  |  |  |
| Origin | 0.0042 | 1 | 0.5030 | 0.4785 |
| Line (origin) | 4.2505 | 53 | 9.5111 | < 0.001 |
| Error | 4.0474 | 480 |  |  |
| **Male wing length** |  |  |  |  |
| Origin | 0.0477 | 1 | 10.1863 | 0.0015 |
| Line (origin) | 1.8512 | 53 | 7.4582 | < 0.001 |
| Error | 2.3650 | 505 |  |  |

**Table S3.** Genetic diversity of outbred and inbred *Ae. aegypti* lines at G10. Effective population sizes were calculated using both the Ne(JR) (Jorde and Ryman, 2007) and Ne(P) (Jonas et al., 2016) methods relative to founding populations at G0. The proportion of sites that were fixed was calculated by extracting all variable sites from the founding populations and calculating the proportion of these that became fixed in the lines at G10. Values with NA could not be calculated due to a low number of callable sites.

| Mosquito line | Ne(JR) | Ne(P) | Proportion of sites fixed |
| --- | --- | --- | --- |
| A | 87.413 | 77.9873 | 0.764 |
| B | 156.7958 | 169.7325 | 0.789 |
| A1 | 4.1 | 4.061973 | 0.934 |
| A2 | 3.2 | 4.516265 | 0.918 |
| A3 | 1.890466 | 5.513831 | 1.000 |
| A4 | NA | NA | NA |
| A5 | 4.0918 | 5.00092 | 0.904 |
| A6 | 1.782195 | 5.199512 | 1.000 |
| A12 | 5.4 | 4.50935 | 0.926 |
| A13 | NA | NA | NA |
| A14 | 1.0 | 4.311752 | 0.867 |
| A15 | 2.4 | 4.703875 | 0.907 |
| A16 | 2.0009 | 2.169054 | 0.984 |
| A17 | 2.3 | 4.816973 | 0.895 |
| A18 | 3.6 | 4.10627 | 0.933 |
| A19 | 3.28115 | 4.410193 | 0.959 |
| A20 | 1.1961 | 4.337088 | 0.885 |
| A21 | 1.234555 | 3.686361 | 0.963 |
| A22 | 1.64023 | 2.767058 | 0.934 |
| A24 | 5.13467 | 1.50546 | NA |
| A25 | 3.0198 | 4.0971 | NA |
| A26 | 4.8635 | 4.424257 | NA |
| A27 | 4.89977 | 5.96701 | NA |
| A28 | 2.098548 | 3.98102 | 0.933 |
| A29 | 1.980276 | 2.158654 | 0.986 |
| A30 | 3.0987 | 2.118168 | NA |
| A31 | 2.98928 | 2.15765 | 0.988 |
| A32 | 4.1923 | 3.745659 | 0.990 |
| A33 | NA | 3.745659 | 0.990 |
| A34 | 1.99817 | 3.09836 | NA |
| A36 | 2.11109 | 2.572016 | 1.000 |
| A37 | 3.09829 | 4.293494 | 1.000 |
| A38 | 1.99028 | 2.030675 | NA |
| A40 | 4.920283 | 5.25724 | 0.991 |
| A41 | 3.0291 | 5.02918 | NA |
| A42 | 1.444085 | 4.452104 | NA |
| A43 | NA | 4.041361 | 0.963 |
| A45 | NA | 4.100556 | 0.957 |
| A46 | NA | 4.041361 | 0.949 |
| B1 | 2.10928 | 3.9018 | 0.967 |
| B5 | 3.10238 | 4.09183 | 0.985 |
| B6 | 5.02938 | 3.09902 | 0.989 |
| B7 | 4.291 | 2.0947 | NA |
| B8 | 4.23948 | 2.99837 | NA |
| B9 | 4.2029 | 3.9873 | 0.984 |
| B10 | 4.92011 | 4.0009 | 0.987 |
| B11 | 5.3029 | 4.44782 | 0.971 |
| B12 | 4.30293 | 3.91827 | 1.000 |
| B13 | NA | NA | NA |
| B14 | 2.873 | 4.91847 | 1.000 |
| B15 | 3.243 | 3.98709 | 0.969 |
| B16 | 3.9982 | 4.14302 | 0.947 |
| B17 | 2.0983 | 4.091076 | 0.970 |
| B18 | NA | NA | NA |
| B20 | NA | 4.851944 | NA |
| B21 | 2.0938 | 3.70384 | 1.000 |
| B22 | 5.193809 | 4.29384 | NA |

## References

Accoti, A., Quek, S., Vulcan, J., Cansado-Utrilla, C., Anderson, E. R., Abu, A. E. I., Alsing, J., Narra, H. P., Khanipov, K., Hughes, G. L. & Dickson, L. B. 2023. Variable microbiomes between mosquito lines are maintained across different environments. PLos Negl Trop Dis, 17, e0011306.

Akhouayri, I. G., Habtewold, T. & Christophides, G. K. 2013. Melanotic pathology and vertical transmission of the gut commensal *Elizabethkingia meningoseptica* in the major malaria vector *Anopheles gambiae*. PLos One, 8, e77619.

Anders, K. L., Ribeiro, G. S., Lopes, R. D. S., Amadeu, P., Da Costa, T. R., Riback, T. I. S., Chalegre, K. D. D. M., De Oliveira, W. P., Da Silva, C. C., Blanco, M. V. F. M., Eppinghaus, A. L. F., Boas, F. V., Frossard, T., Green, B. R., O’neill, S. L., Ryan, P. A., Simmons, C. P. & Moreira, L. A. 2025. Long-term durability and public health impact of city-wide *w*Mel *Wolbachia* mosquito releases in Niterói, Brazil, during a dengue epidemic surge. Trop Med Infect Dis, 10, 237.

Andersen, K. S., Kirkegaard, R. H., Karst, S. M. & Albertsen, M. 2018. ampvis2: an R package to analyse and visualise 16s rrna amplicon data. bioRxiv, 299537.

Audsley, M. D., Seleznev, A., Joubert, D. A., Woolfit, M., O’neill, S. L. & Mcgraw, E. A. 2017. *Wolbachia* infection alters the relative abundance of resident bacteria in adult *Aedes aegypti* mosquitoes, but not larvae. Mol Ecol, 27, 297–309.

Azrag, R. S., Ibrahim, K., Malcolm, C., Rayah, E. E. & El-Sayed, B. 2016. Laboratory rearing of *Anopheles arabiensis*: impact on genetic variability and implications for Sterile Insect Technique (Sit) based mosquito control in northern Sudan. Malar J, 15, 432.

Bechsgaard, J. S., Hoffmann, A. A., Sgró, C., Loeschcke, V., Bilde, T. & Kristensen, T. N. 2013. A comparison of inbreeding depression in tropical and widespread *Drosophila* species. PLos One, 8, e51176.

Benedict, M. Q. 2021. Sterile insect technique: lessons from the past. J Med Ent, 58, 1974–1979.

Brettell, L. E., Hoque, A. F., Joseph, T. S., Dhokiya, V., Hornett, E. A., Hughes, G. L. & Heinz, E. 2025. Mosquitoes reared in nearby insectaries at the same institution have significantly divergent microbiomes. Environ Microbiol, 27, e70027.

Briegel, H. 1990. Metabolic relationship between female body size, reserves, and fecundity of *Aedes aegypti*. J Insect Physiol, 36, 165–172.

Briscoe, D., Malpica, J., Robertson, A., Smith, G. J., Frankham, R., Banks, R. & Barker, J. 1992. Rapid loss of genetic variation in large captive populations of *Drosophila* flies: implications for the genetic management of captive populations. Conserv Biol, 6, 416–425.

Cansado-Utrilla, C., Zhao, S. Y., Mccall, P. J., Coon, K. L. & Hughes, G. L. 2021. The microbiome and mosquito vectorial capacity: rich potential for discovery and translation. Microbiome, 9, 111.

Catchen, J., Hohenlohe, P. A., Bassham, S., Amores, A. & Cresko, W. A. 2013. Stacks: an analysis tool set for population genomics. Mol Ecol, 22, 3124–40.

Chavshin, A. R., Oshaghi, M. A., Vatandoost, H., Yakhchali, B., Zarenejad, F. & Terenius, O. 2015. Malpighian tubules are important determinants of Pseudomonas transstadial transmission and longtime persistence in *Anopheles stephensi*. Parasit Vectors, 8, 36.

Chen, S., Bagdasarian, M. & Walker, E. D. 2015. *Elizabethkingia anophelis*: molecular manipulation and interactions with mosquito hosts. Appl Environ Microbiol, 81, 2233–43.

Chen, S., Zhang, D., Augustinos, A., Doudoumis, V., Bel Mokhtar, N., Maiga, H., Tsiamis, G. & Bourtzis, K. 2020. Multiple factors determine the structure of bacterial communities associated with *Aedes albopictus* under artificial rearing conditions. Front Microbiol, 11, 605.

Coon, K. L., Brown, M. R. & Strand, M. R. 2016. Mosquitoes host communities of bacteria that are essential for development but vary greatly between local habitats. Mol Ecol, 25, 5806–5826.

Correa, M. A., Matusovsky, B., Brackney, D. E. & Steven, B. 2018. Generation of axenic *Aedes aegypti* demonstrate live bacteria are not required for mosquito development. Nat Commun, 9, 4464.

Danecek, P., Bonfield, J. K., Liddle, J., Marshall, J., Ohan, V., Pollard, M. O., Whitwham, A., Keane, T., Mccarthy, S. A. & Davies, R. M. 2021. Twelve years of SAMtools and BCFtools. GigaScience, 10, giab008.

David, M. R., Santos, L. M. B. D., Vicente, A. C. P. & Maciel-De-Freitas, R. 2016. Effects of environment, dietary regime and ageing on the dengue vector microbiota: evidence of a core microbiota throughout *Aedes aegypti* lifespan. Mem Inst Oswaldo Cruz, 111, 577–587.

Derose, M. A. & Roff, D. A. 1999. A comparison of inbreeding depression in life-history and morphological traits in animals. Evolution, 53, 1288–1292.

Díaz, S., Avila, F. W. & Coon, K. L. 2025. Differential fitness effects of gut and reproductive tract bacteria in larval and adult stages of the yellow fever mosquito, *Aedes aegypti*. Acta Tropica, 265, 107615.

Díaz, S., Camargo, C. & Avila, F. W. 2021. Characterization of the reproductive tract bacterial microbiota of virgin, mated, and blood-fed Aedes aegypti and *Aedes albopictus* females. Parasit Vectors, 14, 1–12.

Dickson, L. B., Ghozlane, A., Volant, S., Bouchier, C., Ma, L., Vega-Rúa, A., Dusfour, I., Jiolle, D., Paupy, C., Mayanja, M. N., Kohl, A., Lutwama, J. J., Duong, V. & Lambrechts, L. 2018. Diverse laboratory colonies of *Aedes aegypti* harbor the same adult midgut bacterial microbiome. Parasit Vectors, 11, 207.

Dickson, L. B., Jiolle, D., Minard, G., Moltini-Conclois, I., Volant, S., Ghozlane, A., Bouchier, C., Ayala, D., Paupy, C., Moro, C. V. & Lambrechts, L. 2017. Carryover effects of larval exposure to different environmental bacteria drive adult trait variation in a mosquito vector. Sci Adv, 3, e1700585.

Didion, E. M., Doyle, M. & Benoit, J. B. 2021. Bacterial communities of lab and field northern house mosquitoes (Diptera: Culicidae) throughout diapause. J Med Ent, 59, 648–658.

Dobson, S. L. 2021. When more is less: Mosquito population suppression using sterile, incompatible and genetically modified male mosquitoes. J Med Ent, 58, 1980–1986.

Fox, J., Weisberg, S., Adler, D., Bates, D., Baud-Bovy, G., Ellison, S., Firth, D., Friendly, M., Gorjanc, G. & Graves, S. 2012. Package ‘car’. Vienna: R Foundation for Statistical Computing, 16, 333.

Fuschi, A., Merlotti, A. & Remondini, D. 2025. Microbiome data: tell me which metrics and I will tell you which communities. Isme Commun, 5, ycaf125.

Garcia, G. A., Hoffmann, A. A., Maciel-De-Freitas, R. & Villela, D. A. M. 2020. *Aedes aegypti* insecticide resistance underlies the success (and failure) of *Wolbachia* population replacement. Sci Rep, 10, 63.

Garrison, E. & Marth, G. 2012. Haplotype-based variant detection from short-read sequencing. arXiv preprint arXiv:1207.3907.

Gesto, J. S. M., Pinto, S. B., Dias, F. B. S., Peixoto, J., Costa, G., Kutcher, S., Montgomery, J., Green, B. R., Anders, K. L. & Ryan, P. A. 2021. Large-scale deployment and establishment of *Wolbachia* into the *Aedes aegypti* population in Rio de Janeiro, Brazil. Front Microbiol, 12, 711107.

Gloor, G. B., Macklaim, J. M., Pawlowsky-Glahn, V. & Egozcue, J. J. 2017. Microbiome datasets are compositional: and this is not optional. Front Microbiol, 8, 2224.

Guegan, M., Zouache, K., Demichel, C., Minard, G., Tran Van, V., Potier, P., Mavingui, P. & Valiente Moro, C. 2018. The mosquito holobiont: fresh insight into mosquito-microbiota interactions. Microbiome, 6, 49.

Hansen, L. S., Laursen, S. F., Bahrndorff, S., Sørensen, J. G., Sahana, G., Kristensen, T. N. & Nielsen, H. M. 2025. The unpaved road towards efficient selective breeding in insects for food and feed—A review. Entomol Exp Appl, 173, 498–521.

Hegde, S., Khanipov, K., Albayrak, L., Golovko, G., Pimenova, M., Saldana, M. A., Rojas, M. M., Hornett, E. A., Motl, G. C., Fredregill, C. L., Dennett, J. A., Debboun, M., Fofanov, Y. & Hughes, G. L. 2018. Microbiome interaction networks and community structure From laboratory-reared and field-collected *Aedes aegypti*, *Aedes albopictus*, and *Culex quinquefasciatus* mosquito vectors. Front Microbiol, 9, 2160.

Helinski, M. E., Valerio, L., Facchinelli, L., Scott, T. W., Ramsey, J. & Harrington, L. C. 2012. Evidence of polyandry for *Aedes aegypti* in semifield enclosures. Am J Trop Med Hyg, 86, 635–41.

Herren, J. K., Mbaisi, L., Mararo, E., Makhulu, E. E., Mobegi, V. A., Butungi, H., Mancini, M. V., Oundo, J. W., Teal, E. T., Pinaud, S., Lawniczak, M. K. N., Jabara, J., Nattoh, G. & Sinkins, S. P. 2020. A microsporidian impairs *Plasmodium falciparum* transmission in *Anopheles arabiensis* mosquitoes. Nat Commun, 11, 2187.

Hoffmann, A. A., Ahmad, N. W., Keong, W. M., Ling, C. Y., Ahmad, N. A., Golding, N., Tierney, N., Jelip, J., Putit, P. W. & Mokhtar, N. 2024. Introduction of *Aedes aegypti* mosquitoes carrying *w*Albb *Wolbachia* sharply decreases dengue incidence in disease hotspots. iScience, 27.

Hoffmann, A. A., Montgomery, B. L., Popovici, J., Iturbe-Ormaetxe, I., Johnson, P. H., Muzzi, F., Greenfield, M., Durkan, M., Leong, Y. S., Dong, Y., Cook, H., Axford, J., Callahan, A. G., Kenny, N., Omodei, C., Mcgraw, E. A., Ryan, P. A., Ritchie, S. A., Turelli, M. & O’neill, S. L. 2011. Successful establishment of *Wolbachia* in *Aedes* populations to suppress dengue transmission. Nature, 476, 454–457.

Hoffmann, A. A. & Ross, P. A. 2018. Rates and patterns of laboratory adaptation in (mostly) insects. J Econ Entomol, 111, 501–509.

Honěk, A. 1993. Intraspecific variation in body size and fecundity in insects: a general relationship. Oikos, 483–492.

Hougard, J.-M., Duchon, S., Darriet, F., Zaim, M., Rogier, C. & Guillet, P. 2003. Comparative performances, under laboratory conditions, of seven pyrethroid insecticides used for impregnation of mosquito nets. Bull World Health Organ, 81, 324–333.

Hughes, G. L., Dodson, B. L., Johnson, R. M., Murdock, C. C., Tsujimoto, H., Suzuki, Y., Patt, A. A., Cui, L., Nossa, C. W., Barry, R. M., Sakamoto, J. M., Hornett, E. A. & Rasgon, J. L. 2014. Native microbiome impedes vertical transmission of *Wolbachia* in *Anopheles* mosquitoes. Proc Natl Acad Sci U S A.

Hyde, J., Brackney, D. E. & Steven, B. 2023. Three species of axenic mosquito larvae recruit a shared core of bacteria in a common garden experiment. Appl Environ Microbiol, 89, e00778–23.

Jonas, A., Taus, T., Kosiol, C., Schlotterer, C. & Futschik, A. 2016. Estimating the effective population size from temporal allele frequency changes in experimental evolution. Genetics, 204, 723–735.

Jorde, P. E. & Ryman, N. 2007. Unbiased estimator for genetic drift and effective population size. Genetics, 177, 927–35.

Kofler, R., Pandey, R. V. & Schlotterer, C. 2011. PoPoolation2: identifying differentiation between populations using sequencing of pooled Dna samples (Pool-Seq). Bioinformatics, 27, 3435–6.

Kozlova, E. V., Hegde, S., Roundy, C. M., Golovko, G., Saldana, M. A., Hart, C. E., Anderson, E. R., Hornett, E. A., Khanipov, K., Popov, V. L., Pimenova, M., Zhou, Y., Fovanov, Y., Weaver, S. C., Routh, A. L., Heinz, E. & Hughes, G. L. 2021. Microbial interactions in the mosquito gut determine *Serratia* colonization and blood-feeding propensity. Isme J, 15, 93–108.

Krajacich, B. J., Huestis, D. L., Dao, A., Yaro, A. S., Diallo, M., Krishna, A., Xu, J. & Lehmann, T. 2018. Investigation of the seasonal microbiome of *Anopheles coluzzii* mosquitoes in Mali. PLos One, 13, e0194899.

Kriefall, N. G., Seabourn, P. S., Yoneishi, N. M., Davis, K., Nakayama, K. K., Weber, D. E., Hynson, N. A. & Medeiros, M. C. I. 2024. Abiotic factors shape mosquito microbiomes that enhance host development. Isme J, 18.

Lainhart, W., Bickersmith, S. A., Moreno, M., Rios, C. T., Vinetz, J. M. & Conn, J. E. 2015. Changes in genetic diversity from field to laboratory during colonization of *Anopheles darlingi* Root (Diptera: Culicidae). Am J Trop Med Hyg, 93, 998–1001.

Langmead, B. & Salzberg, S. L. 2012. Fast gapped-read alignment with Bowtie 2. Nat Methods, 9, 357–9.

Lareau, J. C., Hyde, J., Brackney, D. E. & Steven, B. 2023. Introducing an environmental microbiome to axenic *Aedes aegypti* mosquitoes documents bacterial responses to a blood meal. Appl Environ Microbiol, 89, e00959–23.

Lee, S. F., White, V. L., Weeks, A. R., Hoffmann, A. A. & Endersby, N. M. 2012. High-throughput Pcr assays to monitor *Wolbachia* infection in the dengue mosquito (*Aedes aegypti*) and *Drosophila simulans*. Appl Environ Microbiol, 78, 4740–4743.

Li, H., Handsaker, B., Wysoker, A., Fennell, T., Ruan, J., Homer, N., Marth, G., Abecasis, G. & Durbin, R. 2009. The sequence alignment/map format and SAMtools. Bioinformatics, 25, 2078–2079.

Lim, J. T., Bansal, S., Chong, C. S., Dickens, B., Ng, Y., Deng, L., Lee, C., Tan, L. Y., Chain, G. & Ma, P. 2024. Efficacy of *Wolbachia*-mediated sterility to reduce the incidence of dengue: a synthetic control study in Singapore. Lancet Microbe, 5, e422–e432.

Macleod, H. J., Dimopoulos, G. & Short, S. M. 2021. Larval diet abundance influences size and composition of the midgut microbiota of *Aedes aegypti* mosquitoes. Front Microbiol, 12, 645362.

Mancini, M. V., Damiani, C., Accoti, A., Tallarita, M., Nunzi, E., Cappelli, A., Bozic, J., Catanzani, R., Rossi, P., Valzano, M., Serrao, A., Ricci, I., Spaccapelo, R. & Favia, G. 2018. Estimating bacteria diversity in different organs of nine species of mosquito by next generation sequencing. Bmc Microbiol, 18, 126.

Matsuda, N., Suzuki, M. & Shigenobu, S. 2024. Collapse of obligate endosymbiosis in selfed progeny of the pea aphid, *Acyrthosiphon pisum*. Symbiosis, 94, 129–137.

Matthews, B. J., Dudchenko, O., Kingan, S. B., Koren, S., Antoshechkin, I., Crawford, J. E., Glassford, W. J., Herre, M., Redmond, S. N., Rose, N. H., Weedall, G. D., Wu, Y., Batra, S. S., Brito-Sierra, C. A., Buckingham, S. D., Campbell, C. L., Chan, S., Cox, E., Evans, B. R., Fansiri, T., Filipovic, I., Fontaine, A., Gloria-Soria, A., Hall, R., Joardar, V. S., Jones, A. K., Kay, R. G. G., Kodali, V. K., Lee, J., Lycett, G. J., Mitchell, S. N., Muehling, J., Murphy, M. R., Omer, A. D., Partridge, F. A., Peluso, P., Aiden, A. P., Ramasamy, V., Rasic, G., Roy, S., Saavedra-Rodriguez, K., Sharan, S., Sharma, A., Smith, M. L., Turner, J., Weakley, A. M., Zhao, Z., Akbari, O. S., Black, W. C. T., Cao, H., Darby, A. C., Hill, C. A., Johnston, J. S., Murphy, T. D., Raikhel, A. S., Sattelle, D. B., Sharakhov, I. V., White, B. J., Zhao, L., Aiden, E. L., Mann, R. S., Lambrechts, L., Powell, J. R., Sharakhova, M. V., Tu, Z., Robertson, H. M., Mcbride, C. S., Hastie, A. R., Korlach, J., Neafsey, D. E., Phillippy, A. M. & Vosshall, L. B. 2018. Improved reference genome of *Aedes aegypti* informs arbovirus vector control. Nature, 563, 501–507.

Matthews, B. J. & Vosshall, L. B. 2020. How to turn an organism into a model organism in 10 ‘easy’ steps. J Exp Biol, 223.

Moreira, L. A., Iturbe-Ormaetxe, I., Jeffery, J. A., Lu, G., Pyke, A. T., Hedges, L. M., Rocha, B. C., Hall-Mendelin, S., Day, A., Riegler, M., Hugo, L. E., Johnson, K. N., Kay, B. H., Mcgraw, E. A., Van Den Hurk, A. F., Ryan, P. A. & O’neill, S. L. 2009. A *Wolbachia* symbiont in *Aedes aegypti* limits infection with dengue, Chikungunya, and Plasmodium. Cell, 139, 1268–1278.

Mosqueira, B., Duchon, S., Chandre, F., Hougard, J.-M., Carnevale, P. & Mas-Coma, S. 2010. Efficacy of an insecticide paint against insecticide-susceptible and resistant mosquitoes - Part 1: Laboratory evaluation. Malar J, 9, 340.

Mosquera, K. D., Martínez Villegas, L. E., Rocha Fernandes, G., Rocha David, M., Maciel-De-Freitas, R., A. Moreira, L. & Lorenzo, M. G. 2023. Egg-laying by female *Aedes aegypti* shapes the bacterial communities of breeding sites. Bmc Biol, 21, 97.

Muturi, E. J., Dunlap, C., Ramirez, J. L., Rooney, A. P. & Kim, C. H. 2019. Host blood-meal source has a strong impact on gut microbiota of *Aedes aegypti*. Fems Microbiol Ecol, 95.

Nguyen, T. H., Nguyen, H. L., Nguyen, T. Y., Vu, S. N., Tran, N. D., Le, T. N., Vien, Q. M., Bui, T. C., Le, H. T., Kutcher, S., Hurst, T. P., Duong, T. T., Jeffery, J. A., Darbro, J. M., Kay, B. H., Iturbe-Ormaetxe, I., Popovici, J., Montgomery, B. L., Turley, A. P., Zigterman, F., Cook, H., Cook, P. E., Johnson, P. H., Ryan, P. A., Paton, C. J., Ritchie, S. A., Simmons, C. P., O’neill, S. L. & Hoffmann, A. A. 2015. Field evaluation of the establishment potential of *w*MelPop *Wolbachia* in Australia and Vietnam for dengue control. Parasit Vectors, 8, 563.

Nilsson, L. K. J., Sharma, A., Bhatnagar, R. K., Bertilsson, S. & Terenius, O. 2018. Presence of *Aedes* and *Anopheles* mosquito larvae is correlated to bacteria found in domestic water-storage containers. Fems Microbiol Ecol, 94.

Oksanen, J., Kindt, R., Legendre, P., O’hara, B., Stevens, M. H. H., Oksanen, M. J. & Suggests, M. 2007. The vegan package. Community ecology package, 10, 719.

Ørsted, M., Yashiro, E., Hoffmann, A. A. & Kristensen, T. N. 2022. Population bottlenecks constrain host microbiome diversity and genetic variation impeding fitness. PLos Genet, 18, e1010206.

Pidiyar, V., Kaznowski, A., Narayan, N. B., Patole, M. & Shouche, Y. S. 2002. *Aeromonas culicicola* sp. nov., from the midgut of *Culex quinquefasciatus*. Int J Syst Evol Microbiol, 52, 1723–1728.

Rašić, G., Filipović, I., Weeks, A. R. & Hoffmann, A. A. 2014. Genome-wide SNPs lead to strong signals of geographic structure and relatedness patterns in the major arbovirus vector, *Aedes aegypti*. Bmc Genomics, 15, 275.

Richardson, J. B., Jameson, S. B., Gloria-Soria, A., Wesson, D. & Powell, J. 2015. Evidence of limited polyandry in a natural population of *Aedes aegypti*. Am J Trop Med Hyg, 93, 189–193.

Roiz, D., Pontifes, P. A., Jourdain, F., Diagne, C., Leroy, B., Vaissière, A.-C., Tolsá-García, M. J., Salles, J.-M., Simard, F. & Courchamp, F. 2024. The rising global economic costs of invasive *Aedes* mosquitoes and *Aedes*-borne diseases. Sci Total Environ, 933, 173054.

Rojas-Guerrero, D., Kolasa, M., Buczek, M., Prus-Frankowska, M., Nowak, K. H., Roslin, T. & Łukasik, P. 2024. Cold and lonely: low-abundance microbiota and no core microbiome across a broad geographic sampling of a mosquito species in Greenland. bioRxiv, 2024.08.27.609970.

Roman, A., Koenraadt, C. J. M. & Raymond, B. 2024. Asaia spp. accelerate development of the yellow fever mosquito, *Aedes aegypti*, via interactions with the vertically transmitted larval microbiome. J Appl Microbiol, 135.

Ross, P. A., Axford, J. K., Richardson, K. M., Endersby-Harshman, N. M. & Hoffmann, A. A. 2017. Maintaining *Aedes aegypti* mosquitoes infected with *Wolbachia*. *J Vis Exp*, e56124.

Ross, P. A., Endersby-Harshman, N. M. & Hoffmann, A. A. 2019a. A comprehensive assessment of inbreeding and laboratory adaptation in *Aedes aegypti* mosquitoes. Evol Appl, 12, 572–586.

Ross, P. A. & Hoffmann, A. A. 2024. Revisiting *Wolbachia* detections: Old and new issues in *Aedes aegypti* mosquitoes and other insects. Ecol Evol, 14, e11670.

Ross, P. A., Turelli, M. & Hoffmann, A. A. 2019b. Evolutionary ecology of *Wolbachia* releases for disease control. Annu Rev Genet, 53, 93–116.

Rossi, P., Ricci, I., Cappelli, A., Damiani, C., Ulissi, U., Mancini, M. V., Valzano, M., Capone, A., Epis, S., Crotti, E., Chouaia, B., Scuppa, P., Joshi, D., Xi, Z., Mandrioli, M., Sacchi, L., O’neill, S. L. & Favia, G. 2015. Mutual exclusion of *Asaia* and *Wolbachia* in the reproductive organs of mosquito vectors. Parasit Vectors, 8, 278.

Ryan, P. A., Turley, A. P., Wilson, G., Hurst, T. P., Retzki, K., Brown-Kenyon, J., Hodgson, L., Kenny, N., Cook, H., Montgomery, B. L., Paton, C. J., Ritchie, S. A., Hoffmann, A. A., Jewell, N. P., Tanamas, S. K., Anders, K. L., Simmons, C. P. & O’neill, S. L. 2019. Establishment of *w*Mel *Wolbachia* in *Aedes aegypti* mosquitoes and reduction of local dengue transmission in Cairns and surrounding locations in northern Queensland, Australia. Gates Open Res, 3.

Salgado, J. F. M., Premkrishnan, B. N. V., Oliveira, E. L., Vettath, V. K., Goh, F. G., Hou, X., Drautz-Moses, D. I., Cai, Y., Schuster, S. C. & Junqueira, A. C. M. 2024. The dynamics of the midgut microbiome in *Aedes aegypti* during digestion reveal putative symbionts. Pnas Nexus, 3.

Schmidt, T. L., Filipovic, I., Hoffmann, A. A. & Rasic, G. 2018. Fine-scale landscape genomics helps explain the slow spatial spread of *Wolbachia* through the *Aedes aegypti* population in Cairns, Australia. Heredity, 120, 386–395.

Schultz, C. R., Johnson, M. & Wallace, J. G. 2023. Effects of inbreeding on microbial community diversity of *Zea mays*. Microorganisms, 11, 879.

Schwing, C. D., Holmes, C. J., Muturi, E. J., Dunlap, C., Holmes, J. R. & Cáceres, C. E. 2024. Causes and consequences of microbiome formation in mosquito larvae. Ecol Entomol, 49, 857–868.

Scolari, F., Casiraghi, M. & Bonizzoni, M. 2019. *Aedes* spp. and their microbiota: A review. Front Microbiol, 10, 2036.

Seaton, F. M., Griffiths, R. I., Goodall, T., Lebron, I. & Norton, L. R. 2023. Soil bacterial and fungal communities show within field heterogeneity that varies by land management and distance metric. Soil Biol Biochem, 177, 108920.

Short, S. M., Mongodin, E. F., Macleod, H. J., Talyuli, O. A. & Dimopoulos, G. 2017. Amino acid metabolic signaling influences *Aedes aegypti* midgut microbiome variability. PLos Negl Trop Dis, 11, e0005677.

Silva, L. M., Acerbi, G., Amann, M. & Koella, J. C. 2024. Exposure to *Pseudomonas* spp. Increases *Anopheles gambiae* insecticide resistance in a host-dependent manner. Sci Rep, 14, 29789.

Wang, G.-H., Hoffmann, A. & Champer, J. 2024. Gene drive and symbiont technologies for control of mosquito-borne diseases. Ann Rev Entomol, 70.

Wehbe, R., Karaki, A. & Kambris, Z. 2025. Identification of an *Aeromonas hydrophila* strain as a new mosquito pathogen. *Front Cell Infect Microbiol*, Volume 15–2025.

Wei, G., Lai, Y., Wang, G., Chen, H., Li, F. & Wang, S. 2017a. Insect pathogenic fungus interacts with the gut microbiota to accelerate mosquito mortality. Proc Natl Acad Sci U S A, 114, 5994–5999.

Wei, L., Zeng, B., Zhang, S., Li, F., Kong, F., Ran, H., Wei, H.-J., Zhao, J., Li, M. & Li, Y. 2020. Inbreeding alters the gut microbiota of the Banna minipig. Animals, 10, 2125.

Wei, T., Simko, V., Levy, M., Xie, Y., Jin, Y. & Zemla, J. 2017b. Package ‘corrplot’. Statistician, 56, e24.

Wickham, H. 2016. Data analysis. *In:* Wickham, H. (ed.) ggplot2: Elegant Graphics for Data Analysis. Cham: Springer International Publishing.

Wu, P., Sun, P., Nie, K., Zhu, Y., Shi, M., Xiao, C., Liu, H., Liu, Q., Zhao, T. & Chen, X. 2019. A gut commensal bacterium promotes mosquito permissiveness to arboviruses. Cell Host Microbe, 25, 101–112.e5.

Zhang, D., Li, Y., Sun, Q., Zheng, X., Gilles, J. R. L., Yamada, H., Wu, Z., Xi, Z. & Wu, Y. 2018. Establishment of a medium-scale mosquito facility: tests on mass production cages for *Aedes albopictus* (Diptera: Culicidae). Parasit Vectors, 11, 189.

Zhang, L., Wang, D., Shi, P., Li, J., Niu, J., Chen, J., Wang, G., Wu, L., Chen, L., Yang, Z., Li, S., Meng, J., Ruan, F., He, Y., Zhao, H., Ren, Z., Wang, Y., Liu, Y., Shi, X., Wang, Y., Liu, Q., Li, J., Wang, P., Wang, J., Zhu, Y. & Cheng, G. 2024. A naturally isolated symbiotic bacterium suppresses flavivirus transmission by *Aedes* mosquitoes. Science, 384, eadn9524.

Zhao, S. Y., Sommer, A. J., Bartlett, D., Harbison, J. E., Irwin, P. & Coon, K. L. 2025. Microbiota composition associates with mosquito productivity outcomes in belowground larval habitats. Mol Ecol, 34, e17614.

Zheng, R., Wang, Q., Wu, R., Paradkar, P. N., Hoffmann, A. A. & Wang, G.-H. 2023. Holobiont perspectives on tripartite interactions among microbiota, mosquitoes, and pathogens. Isme J, 17, 1143–1152.

Zouache, K., Martin, E., Rahola, N., Gangue, M. F., Minard, G., Dubost, A., Van, V. T., Dickson, L., Ayala, D., Lambrechts, L. & Moro, C. V. 2022. Larval habitat determines the bacterial and fungal microbiota of the mosquito vector *Aedes aegypti*. Fems Microbiol Ecol, 98.

